# Distinct Expansion of Group II Introns Depends on the Type of Intron-encoded Protein and Genomic Signatures in Prokaryotes

**DOI:** 10.1101/2021.09.28.462093

**Authors:** Masahiro C. Miura, Shohei Nagata, Satoshi Tamaki, Masaru Tomita, Akio Kanai

## Abstract

Group II introns (G2Is) are self-splicing ribozymes that have retroelement characteristics in prokaryotes. Although G2Is are considered an important factor in the evolution of prokaryotes, comprehensive analyses of these introns among the tens of thousands of prokaryotic genomes currently available are still limited. Here, we developed a bioinformatic pipeline that systematically collects G2Is and applied it to prokaryotic genomes. We found that in bacteria, 25% (447 of 1,790) of the total representative species had an average of 5.3 G2Is, and in archaea, 9% (28 of 296) of the total representative species had an average of 3.0 G2Is. The greatest number of G2Is per species was 101 in *Arthrospira platensis* (phylum Cyanobacteriota). A comprehensive sequence analysis of the intron-encoded protein (IEP) in each G2I sequence was conducted and resulted in the addition of three new IEP classes (U1–U3) to the previous classification. This analysis suggested that about 30% of all IEPs are noncanonical IEPs. The number of G2Is per species was defined almost at the phylum level, and the type of IEP was associated as a factor in the G2I increase, i.e. there was an explosive increase in G2Is with bacterial C-type IEPs in the phylum Firmicutes and in G2Is with CL-type IEPs in the phylum Cyanobacteriota. We also systematically analyzed the relationship between genomic signatures and the mechanism of these increases in G2Is. This is the first study to systematically characterize G2Is in the prokaryotic phylogenies.

## Introduction

Introns are present in various forms across the three domains of life, bacteria, archaea, and eukaryota. The best known is the pre-mRNA intron in the eukaryotic nuclear genome, which is spliced by spliceosomes (Galej et al. 2018). Many self-splicing introns, such as group I introns (G1I) and group II introns (G2I), have been found in prokaryotes (both bacteria and archaea) and eukaryotic organelles (Edgell et al. 2011). Bulge–helix–bulge (BHB) introns are also inserted into some pre-tRNAs and pre-rRNAs in archaea, and into some pre-tRNAs in eukaryotes, and are enzymatically spliced by endoribonucleases and RNA ligases (Tang et al. 2002; Yoshihisa 2014). We developed a computational program to comprehensively extract tRNA introns from predominantly archaeal genomes and detected tRNA genes that are disrupted in various ways (Sugahara et al. 2008; Fujishima et al. 2009). We reported that among the archaeal tRNA introns, there are mobile tRNA introns in the archaeal order Thermoproteales (Fujishima et al. 2010). We also discovered a novel endoribonuclease involved in the excision of pre-tRNA introns and reported its coevolution with a specific type of pre-tRNA intron (Fujishima et al. 2011). In the context of this research, and based on the facts that the tertiary structure of spliceosome is similar to that of G2Is (indicating an evolutionary relationship between the spliceosome and G2I) and that a huge number of complete prokaryotic genome sequences is currently available, we decided to systematically analyze the G2I, which comprises an intron, a ribozyme, and a mobile element in prokaryotes (Lambowitz and Belfort 2015).

Standard G2I RNAs are divided structurally into domains I–VI based on their RNA secondary structures. Some of these secondary-structural domains have special functions. For example, the domain I region contains the target-binding sites for G2I transposition, and the domain V region acts as the center of ribozyme activity in self-splicing (Lambowitz and Belfort 2015). Although G2I is a ribozyme, intron-encoded proteins (IEPs) are encoded in domain IV (Michel et al. 1982). Each IEP consist of a reverse transcriptase (RT) domain and other domains (e.g. DNA-binding domain and endonuclease [En] domain). In addition to its reverse transcriptase activity, the RT domain functions as a maturase, which binds to the G2I RNA and stabilizes its tertiary structure. During transposition, the DNA-binding domain and En domain bind to the target region of DNA and break the bottom DNA strand (the DNA strand into which G2I is not integrated), respectively. However, in some types of IEP, the En domain is not structurally included in the IEP and is not essential for transposition. Thus, the G2I RNA and IEP form a complex, and the self-splicing and transposition activities are conducted by the cooperative action of each functional domain of the G2I RNA and IEP. Interestingly, it has been suggested that the G2I ribonucleoprotein (RNP) and spliceosomes, which are giant RNPs responsible for splicing eukaryotic pre-mRNA, have a common evolutionary origin because their tertiary structures and the basic processes of their splicing reactions are very similar (Zimmerly and Semper 2015; Novikova and Belfort 2017). G2Is are classified based on the similarity of the secondary structures of their RNAs and the similarity of the amino acid sequences of their IEPs (Michel et al. 1989; Zimmerly et al. 2001), and there are differences in the self-splicing reaction pathways and the target regions of the retromobility reactions between the classified groups (McNeil et al. 2016). Some G2Is do not have an IEP (open reading frame [ORF]-less type) (Simon et al. 2008). In rare cases, nonstandard G2Is have also been reported in which the RNA secondary structure is degenerate, the protein encoded by the ORF is largely lacking due to frameshift, or the encoded protein is a homing endonuclease. However, at least some of the nonstandard G2Is have splicing activity (Salman et al. 2012; McNeil et al. 2014).

Attempts have been made to computationally identify regions corresponding to G2Is in genomic DNA data. In the early 2000s, Zimmerly collected bacterial and archaeal G2Is using a sequence-similarity-search-based method and published it as a database (Zimmerly et al. 2001; Dai and Zimmerly 2002, 2003). That research group also collected diverse sequences by creating a descriptor of RNA secondary structures (Simon et al. 2008). Since then, the G2I sequence published by that group has been used as a query by other groups, and G2Is have been collected continuously from genomic sequences (Leclercq et al. 2011; Abebe et al. 2013; Toro and Nisa-Martinez 2014; Titov et al. 2019). These analyses have shown the diversity of G2Is, the numbers of G2Is per species, and their insertion positions in genomes. Toro and Nisa-Martínez collected various reverse transcriptases, including the IEPs in G2Is, and showed that the G2Is can be classified into 13 types based on the similarities of their RT domains (Toro and Nisa-Martinez 2014). It has also been shown that in most bacteria, the number of G2Is per genome is ≤ 3 and rarely exceeds 10 (Leclercq et al. 2011; Waldern et al. 2020). Moreover, G2Is often nest within the intergenic regions that are not harmful to the host bacterium or within other mobile genetic elements, including G2Is or related genes, whereas some G2Is have been reported to disrupt essential genes (Novikova et al. 2014; Waldern et al. 2020).

As mentioned above, numerous studies have provided much insight into G2Is with bioinformatic analyses. However, while the number of prokaryote genomes registered in public databases is continuing to increase, the bioinformatic analyses to address the overall picture of the G2Is in the database are still limited. Compared with bacteria, archaeal G2Is are rarely the subject of comprehensive analyses and there are few reports of their distribution on the phylogenetic tree. Fragmentary sequences of G2Is also exist in the genomes of prokaryotes, but these noncanonical sequences have not been the subject of analysis in most cases. Furthermore, most analyses have focused on G2I itself, and few studies have comprehensively investigated the correspondence among G2I characteristics, host taxa, and genomic sequence signatures. Therefore, we attempted to clarify these points by developing a bioinformatic pipeline that systematically collects G2Is and applied it to the complete prokaryotic genomes (approximately 15,000) currently available. By classifying the collected G2Is, we clarified the possible existence of new G2I groups based on the IEP sequences, and the spread of fragmented noncanonical IEPs in prokaryotes. The phylogenetic information on the collected G2Is and the host genomes allowed us to establish the detailed distributions of G2Is in bacteria and archaea according to their IEP types. The number of G2Is was generally defined at the phylum level, but there were many cases in which the numbers differed significantly among closely related species. A dramatic increase in G2Is in prokaryotes occurred with combinations of specific IEP types and bacterial taxa, and these increases may be associated with particular genomic signatures, such as transcription terminators and GC skew. All the G2I data analyzed in this study are summarized in supplementary files, which we hope will be useful for genome-scale studies of prokaryotic transposable elements.

## Results and Discussion

### Comprehensive Extraction of G2Is from Prokaryotic Genomes

In an attempt to understand the exact distribution of G2Is in prokaryotic genomes, it is necessary to comprehensively identify the genomic region of each G2I. We first constructed a new bioinformatic pipeline that comprehensively identifies G2Is in prokaryotic genomes. This program allowed us to identify genomic sequences containing the most conserved RNA secondary structures of G2Is, domains V and VI, and at least parts of domains I–IV. Another feature of the pipeline is that it can handle G2Is with or without IEPs. A summary of the pipeline is given below (Supplementary Figure S1A). In step 1, the three major domains of the G2Is were extracted: (i) the RT domain of the IEP, (ii) RNA domains V and VI, and (iii) RNA domains I–IV. The RNA secondary structure model registered in the Rfam database was used to extract domains V and VI, and parameters were established to identify 341 of the 347 (98%) G2Is classified as “Eubacterial”, “Archaeal”, or “ORF-less” in the Database for Bacterial Group II Introns (Zbase) (Candales et al. 2012). In step 2, when the RT domain occurred within the 1,300 bases of domain V, the sequence was interpreted as the IEP-containing type of G2I (Supplementary Figure S1B). In step 3, if an RT domain was not considered to be part of G2I in step 2, and domains I–IV were within the 1,300 bases of domain V, the sequence was interpreted as a no-IEP-type or ORF-less G2I (Supplementary Figure S1C). Therefore, when setting the threshold for the distances between domains in steps 2 and 3, we calculated the length of each region of the G2Is in Zbase and in a previous study (Toro and Nisa-Martinez 2014). The threshold was set to 1,300 bases because the distance between the RT domain and domain V was ≤ 1,300 bases in 99% of the identified G2Is. For the ORF-less-type G2Is, the threshold was set to 1,300 bases based on the same criterion.

To verify the pipeline, 12 species with G2Is and 18 species without G2Is in prokaryotes were selected (species with G2Is were selected from those registered in Zbase), and the program to extract G2Is was applied to these genomes (Supplementary Table S1). The results indicated that in 18 species for which no G2I was reported, our program detected no G2Is, i.e., there were no false positive results. Furthermore, our program detected 98% of the G2Is with IEPs and the ORF-less-type G2Is in which no RNA secondary-structural domain was deleted, as shown in Supplementary Table S1. These two types of G2Is accounted for 80% of the 434 G2Is of prokaryotes registered in Zbase, and we consider that the majority of G2Is in prokaryotic genomes belong to one or other of these types. Similarly, most previous studies that collected G2Is from genomic data targeted either one of these two types. This demonstrates that our program can effectively collect more data than was achieved in previous studies. For example, *Sinorhizobium meliloti* has six G2Is with IEPs and *Paraburkholderia xenovorans* has two G2Is with IEPs and one ORF-less G2I, and all of these G2Is were detected with our program. However, in the following cases, there was a slight discrepancy between the results of our program and the prior studies. First, if multiple G2Is had a nested structure, called a ‘twintron’ (Nakamura et al. 2002; Pfreundt and Hess 2015), our program could not detect the outer G2I. This situation was found in the genome of *Thermosynechococcus elongatus* and *Wolbachia* endosymbiont (false negative case #1). Additionally, our program did not detect the G2I-like sequences in *T. elongatus* that lacked IEPs and most of domains I–IV (false negative case #2). Although the program detected G2Is that are considered to be pseudogenes because the IEPs have frameshift mutations, these sequences were not registered in Zbase (false positive case #1). This result was classified as a false positive here, but as explained later, it is classified as a G2I with a noncanonical IEP in this paper. This situation was detected in the genomes of *Enterococcus faecalis* and *Methanosarcina mazei*. In *Methanococcoides burtonii*, a region containing domains V and VI was duplicated before and after the IEP in all four G2Is. This resulted in the presence of two genes per G2I (false positive case #2). We undertook a large-scale data analysis of the main G2Is mentioned above, including the G2I-like sequences but excluding the G2Is with exceptional structures.

### Number of G2Is is Usually Dictated by the Host Phylum, but Can Differ Even Within the Same Phylum

Using our program, we searched for G2Is in 14,506 bacterial genomes and 296 archaeal genomes downloaded from the National Center for Biotechnology Information (NCBI) reference sequence (RefSeq) database. In this way, 12,217 G2I sequences with IEPs and 908 G2I sequences without IEPs were identified. We then analyzed the phylogenetic distribution of the species with G2Is. In bacteria, the results were narrowed to 1,790 representative species in order to eliminate the bias of species registered in the database, and in archaea, the results for all species were included to ensure as many species as possible. As shown in Table 1A, a total of 2,381 G2Is were detected in 447 bacterial species, or approximately 25% of the 1,790 species. Of these, 2,100 were classified as G2Is with IEPs. In previous studies, the details of the method used and the types of G2I targeted differed between research groups, and the abundance of G2Is in bacteria has been reported to be 12%–35% (Leclercq et al. 2011; Candales et al. 2012; Waldern et al. 2020). A previous study claimed that the prevalence of G2Is in bacteria is approximately 25% of the total species (Candales et al. 2012), and our results support this observation. We also confirmed that 29 of the 41 bacterial phyla examined contained G2Is, and that G2Is are widely distributed in bacteria. At the phylum level, Firmicutes B, Cyanobacteriota, Desulfuromonadota, and Myxococcota each contained ≥ 10 species in the dataset, and more than 50% of these species had G2Is. By contrast some phyla had no G2Is among any of the species included in the dataset, such as Deinococcota with 20 species in the dataset. However, among the 447 species containing G2Is, 28 contained only ORF-less-type G2Is. In archaea, our search found a total of 84 G2Is in 28 species, which accounted for approximately 9% of the total archaeal species (Table 1B). At the phylum level, 23% of the species in Halobacterota contained G2Is, whereas fewer than 1% of the species in Crenarchaeota and Euryarchaeota contained G2Is. These results indicate that G2Is exist only in limited archaeal species, in contrast to the spread of G2Is in bacteria. We note that the current analysis was limited to the near-complete prokaryotic genomes in the database, so Candidate Phyla Radiation (CPR) bacteria and Asgard archaea have not been analyzed, while G2Is have also been found in these species in previous studies (Brown et al. 2015; Vosseberg and Snel 2017). The species in which G2Is were found and their locations in these genomes are summarized in Supplementary Table S2. Information on the 14,506 bacterial genomes and 296 archaeal genomes used in this study is summarized in Supplementary Table S3.

**Table 1.**
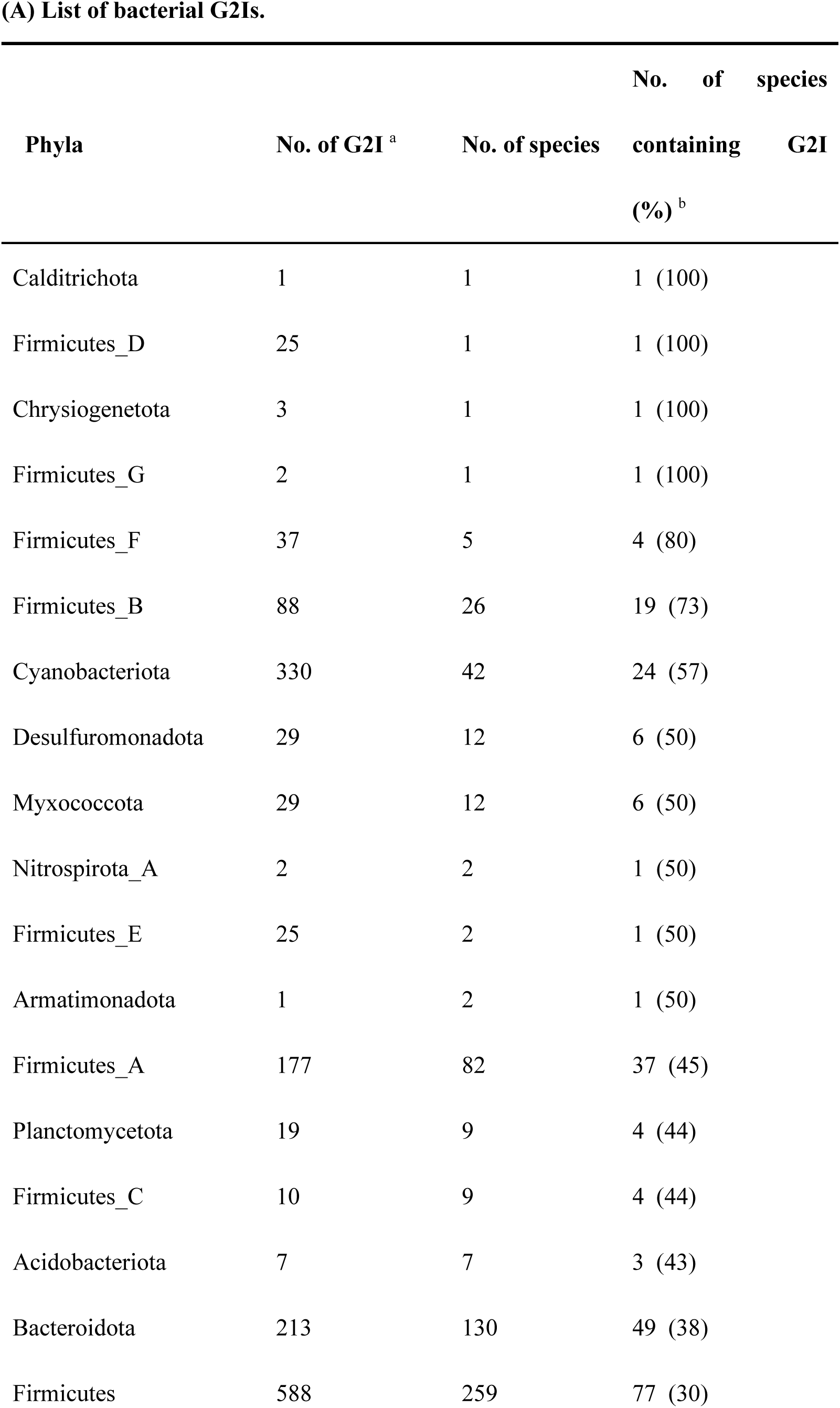
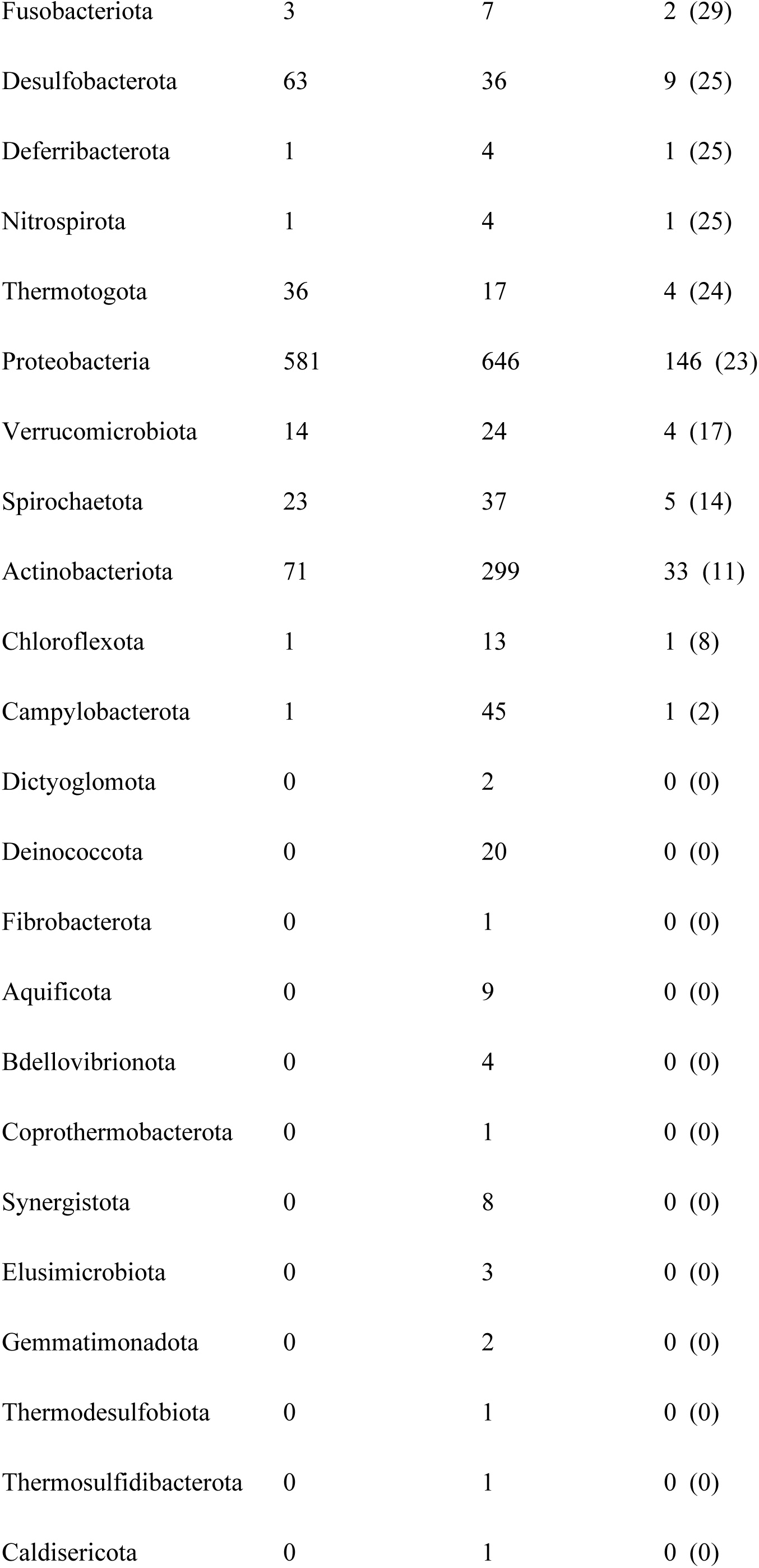
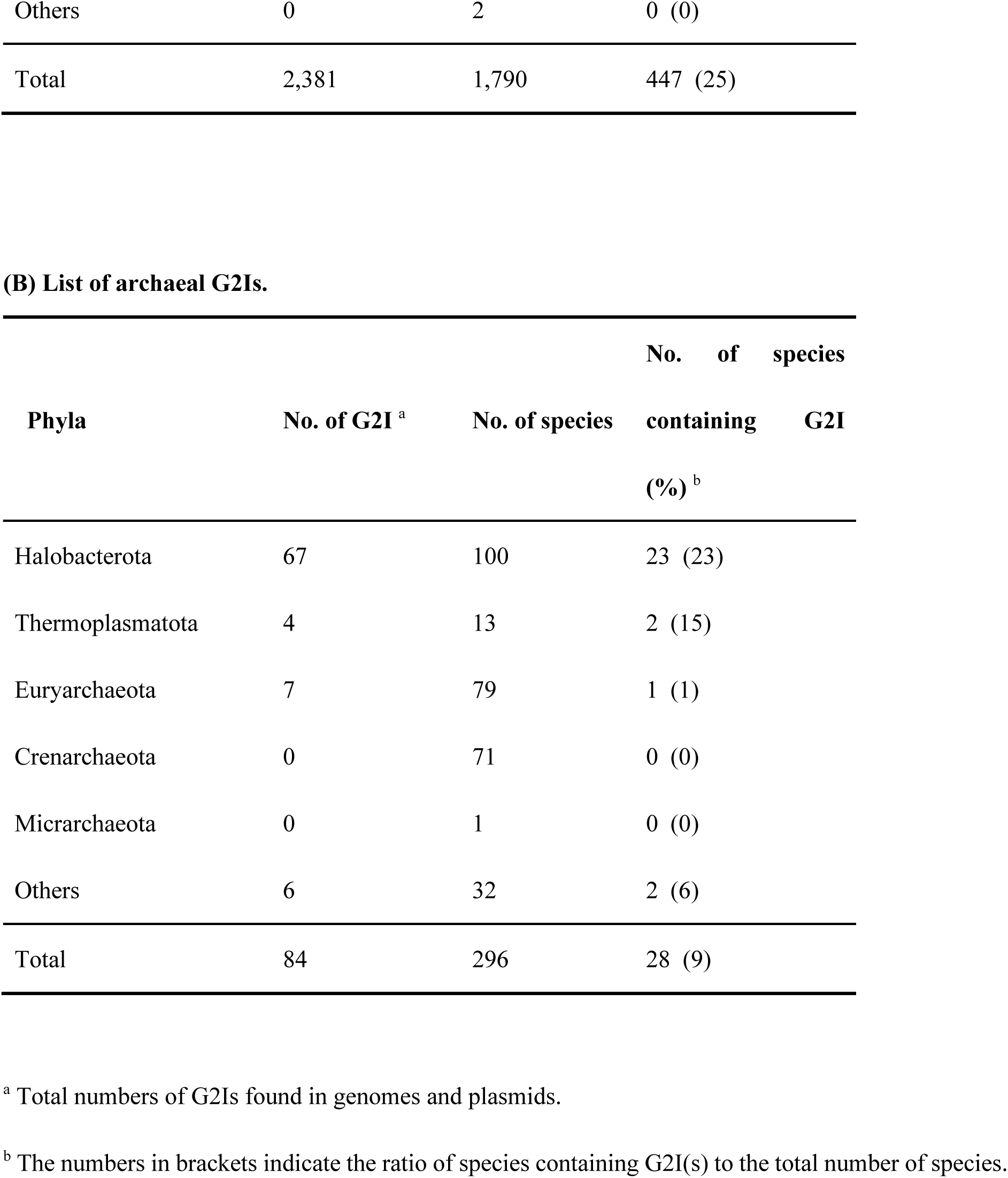
Number of G2Is detected in this study.

We next analyzed the number of G2Is per bacterial species (Figure 1). In the 447 bacterial species containing G2Is, the average number ± standard deviation of G2Is was 5.3 ± 8.8, and the median was 2, suggesting considerable variability. The largest number of G2Is was in *Arthrospira platensis* (RefSeq assembly accession: GCF_000210375.1), which has 101 G2Is. Fifty-four species, or about 12% of the total number of species, had ≥ 10 G2Is. In Actinobacteriota and Campylobacterota, most species had very few G2Is, and few species had ≥ 10 G2Is. Firmicutes B and Cyanobacteriota are phyla in which many species contain G2Is, and 11% and 16% of their species, respectively, had ≥ 10 G2Is. Cyanobacteriota contained three species, *Thermosynechococcus elongatus*, *Trichodesmium erythraeum*, and *Arthrospira platensis*, that reportedly have abundant G2Is (Nakamura et al. 2002; Pfreundt et al. 2014; Xu et al. 2016). We also confirmed the species with very large numbers of G2Is (≥ 50) in Firmicutes and Cyanobacteriota. Based on these results, we infer that the distribution of G2Is differs among phyla, and that the numbers of introns increased rapidly in specific bacterial phyla. We also found that in phyla containing many species, such as Firmicutes and Proteobacteria, there are two groups based on the number of G2Is: the number of G2Is has increased in one group, but they are almost absent in the other group. We also analyzed the relationship between the number of G2Is and the genome size as a factor that might explains the differences in the species distributions and the numbers of G2Is across species of bacteria and archaea. The results showed that the correlation coefficient between the number of G2Is and genome size was low (R^2^ = 0.003 or 0.002), so no relationship was detected (Supplementary Figure S2A). The possibility of a relationship between the number of G2Is and genome size was also examined for each bacterial phylum, but no significant relationship was found in any phylum (Supplementary Figure S2B).

**Fig. 1.**
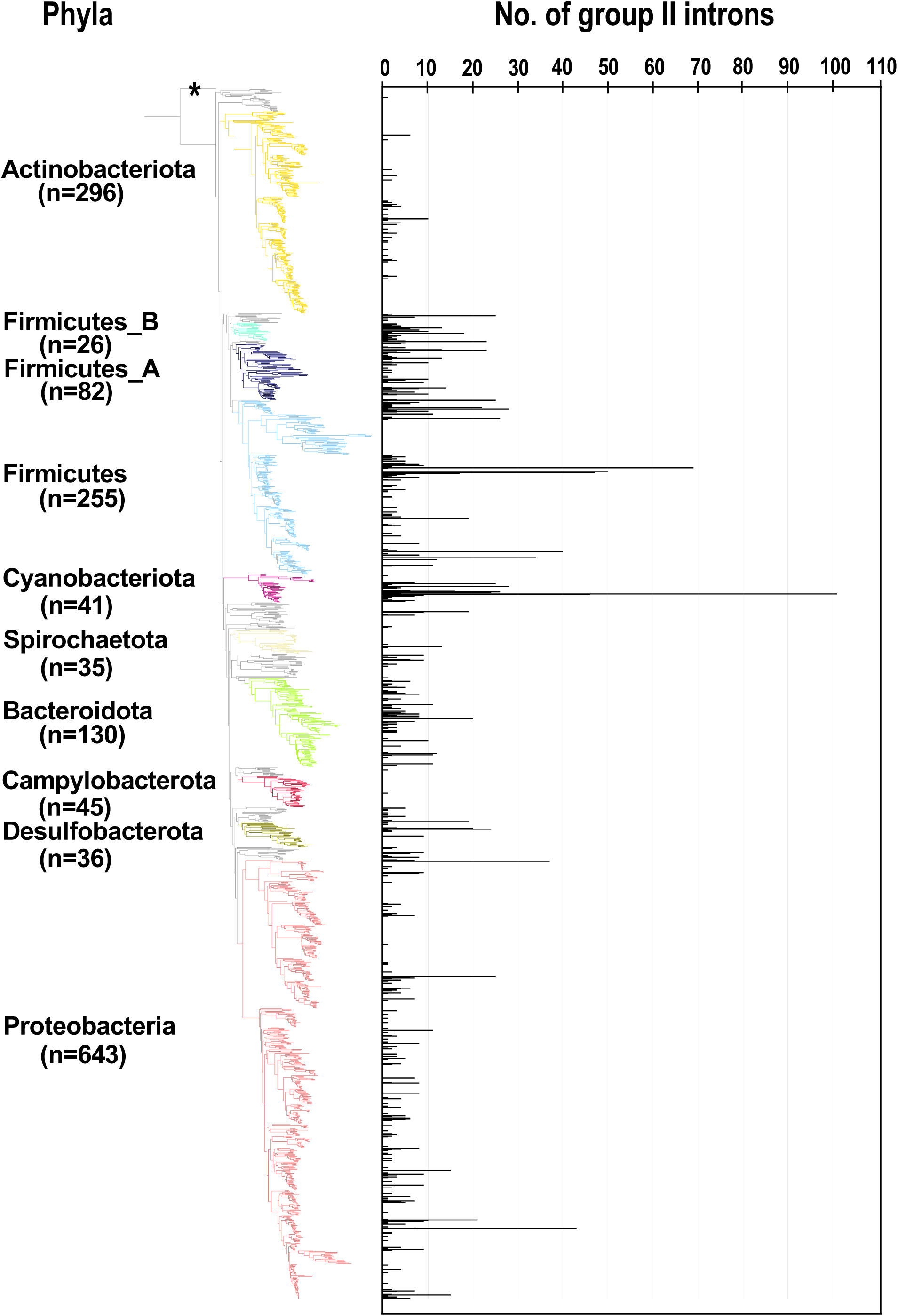
Increase in the number of G2Is in specific bacterial species belonging to each bacterial phylum. The numbers of G2Is in representative complete bacterial genomes (1,775 species) are shown. Bacterial phyla are shown on the left and each corresponding branch on the bacterial phylogenic tree is colored. The numbers in bracket represents the number of species in each phylum. The position of the outgroup (*Candidatus Saccharibacteria* oral taxon TM7x [RefSeq assembly accession: GCF_000803625.1]) is indicated by the asterisk.

### G2Is Present in Large Numbers Have Specific IEP Types

To clarify why the number of G2Is per genome differs for each taxon, we undertook an analysis of the types of IEPs present in the G2Is. First, 12,217 IEPs were extracted from 14,506 bacterial and 296 archaeal genomes (see Materials and Methods). Figure 2A shows a phylogenetic tree constructed from 1,977 sequences of representative IEPs obtained with CD-HIT from 12,217 IEPs. Among the IEPs we collected, there were sequences that were interrupted by a stop codon and sequences that were clearly lacking some functional domains. These incomplete sequences accounted for about 30% of the total 1,977 IEP sequences used to construct the phylogenetic tree (Supplementary Figure S3). Regardless of the species or type of IEP, these “noncanonical” IEPs were widely distributed in the bacterial phylogeny (Supplementary Figures S4 & S5). In contrast, in some cases, because nucleotide sequences similar to the corresponding region of each canonical IEP occur after the stop codon, it is possible that some noncanonical IEP sequences encode complete IEPs, expressed by read-through or frameshift mechanisms. The types of IEPs classified on this phylogenetic tree well reflect the IEP types in previous studies (Toro and Martinez-Abarca 2013; Toro and Nisa-Martinez 2014). We also found that the IEP sequences that were considered “unclassified” in the previous study clustered at three phylogenetic positions, and designated them U1, U2, and U3 on the current phylogenetic tree: U1 is located between the bacterial-g2–g5 clades and bacterial-g6 clade; U2 is located in the same clade position as bacterial-g2–g5, but its distribution differs; and U3 forms a clade in a region located near the bacterial-A type IEPs. U1, U2, and U3 are not monophyletic, but each has different sequence characteristics from the neighboring IEPs on the phylogenetic tree. For example, compared with the amino acid sequences of the bacterial-g2–g5-type IEPs, most of the U1- and U2-type IEPs lack part of the RT domain (Supplementary Figure S6). Compared with the bacterial-A-type IEPs, U3 tends to lack a part of the RT domain and the DNA-binding domain is elongated (Supplementary Figure S7). These results suggest that the U1–U3 lineages constitute new phylogenetic subtypes, at least.

**Fig. 2.**
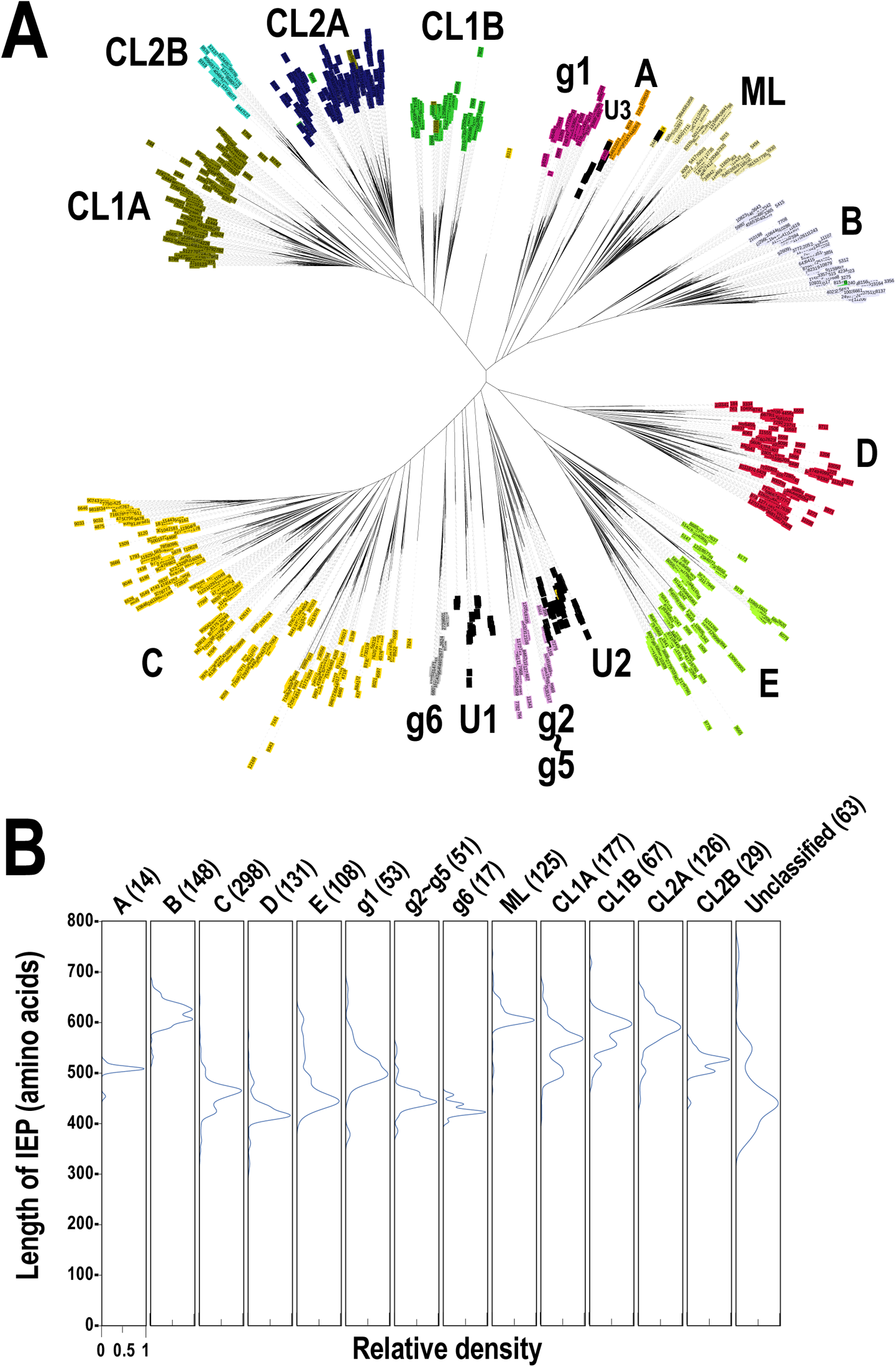
Phylogeny of the prokaryotic IEPs detected in this study. (A) An unrooted phylogenetic tree of representative IEP sets (1,977 proteins) in bacterial or archaeal G2Is. The representative IEP sets were selected based on a similarity analysis (see Materials and Methods, Prediction of IEP Sequences). The types of IEP are: A (bacterial-A, orange), B (bacterial-B, lavender), C (bacterial-C, yellow), D (bacterial-D, red), E (bacterial-E, light green), g1 (bacterial-g1, violet), g2–g5 (bacterial-g2–g5, plum), g6 (bacterial-g6, silver), ML (light yellow), CL1A (olive), CL1B (green), CL2A (blue), and CL2B (turquoise). U1–U3 (black) are newly identified clusters that were previously annotated as “unclassified.” (B) Distribution of the amino acid lengths of each canonical IEP. The peak relative density is set as 1.0 in each case.

Figure 2B shows the distribution of the sequence lengths of the canonical ORFs for each IEP lineage group. Note that because each ORF sequence was selected to be as long as possible when the ORF was extracted, the methionine codon upstream from the exact translation start position can be used, and the sequence length may be overestimated. Most IEP groups had sequence lengths of 400–700 amino acids. Toro et al. reported that the sequence lengths of the RT domains of G2Is range from 240 to 330 amino acids (Toro and Nisa-Martinez 2014). In contrast, we calculated that in regions other than the RT domain, the sequence length varies widely, ranging from 140 to 360 amino acids, depending on the IEP type. This is because some types of IEPs, such as bacterial-B, ML, CL1A, CL1B, and CL2A, are long (peaking near 600 amino acids) because they contain an En domain (Supplementary Figure S1B) (Zimmerly and Semper 2015). In other IEPs without an En domain, the peak length is approximately 400–500 amino acids. Furthermore, although the CL2B-type IEPs have an En domain, the overall length is slightly reduced because the region of the DNA-binding domain is shorter than that in other CL-type IEPs. The unclassified IEP types have two peak lengths, near 550 and 450 amino acids. This is because the three types have different length distributions: U1 is about 350–500 amino acids, U2 is about 350–750 amino acids, and U3 is about 500–550 amino acids. When the distribution of the lengths of the ORFs of noncanonical IEPs is considered, many IEP types have a wide range of length and some very short sequences of about 150–350 amino acids. These are fragmentary IEPs, in which at least part of some domain(s) is not present (Supplementary Figure S8A). By contrast, among noncanonical IEPs, there are several sequences with the same lengths as canonical IEPs. These are sequences in which a relatively long ORF occurs due to a shift in the reading frame, although the same domain structure is not maintained. When the distribution of sequence lengths was analyzed after the canonical and noncanonical IEPs were combined, the distribution of canonical IEPs (Figure 2B) was almost similar to that of combined IEPs, and the short IEPs found among the noncanonical IEPs constituted only a small part of the whole set (Supplementary Figure S8B). We are the first to undertake a comprehensive analysis of noncanonical IEPs in this way.

Next, the distribution of IEP types was examined in the 443 representative bacterial species with G2Is summarized in Supplementary Table S1A, excluding four species not used in the phylogenetic tree (Figure 3). The IEP types corresponded well to specific bacterial phyla, such as bacterial B to Firmicutes, g1 to Actinobacteriota, CL2A to Cyanobacteriota, and CL2B to Cyanobacteriota, whereas bacterial-C, -D, and -E and CL1A are widely distributed throughout the bacterial phyla. The number of G2Is with bacterial-C-type IEPs was high in many species. This result is consistent with a previous study in which more than half of the IEPs found in 1,435 bacteria were of the bacterial-C type (Waldern et al. 2020). Number of G2Is by IEP type in each species is summarized in Supplementary Table S4. To clarify the relationship between the type of IEP and the number of G2Is at the species level, 25 species with ≥ 20 G2Is were selected. Supplementary Figure S9 shows that species belonging to Firmicutes tended to encode bacterial-C-type IEPs, and species belonging to Cyanobacteriota tended to encode CL-type IEPs. Furthermore, the number of G2Is with bacterial-C-type IEPs increased to ≥ 20 in phyla other than Firmicutes, including Bacteroidota, Desulfobacterota, and Proteobacteria. However, CL-type G2Is were significantly increased only in the species belonging to Cyanobacteriota, except for one species, *Vibrio campbellii_A*. A few species (e.g., *Orientia tsutsugamushi* and *Halobacteroides halobius*) had large numbers of bacterial-E-type and unclassified-type G2Is. The number of ORF-less-type G2Is was increased in a wide range of species belonging to Firmicutes, Cyanobacteriota, Desulfobacteriota, and Proteobacteria. As characteristics of host bacteria in which G2Is increase, it has been reported that an obligate endosymbiotic bacterium, which is difficult to culture alone, and a thermophilic bacterium have a large number of G2Is (Mohr et al. 2010; Leclercq et al. 2011; Mohr et al. 2018). In Supplementary Figure S9, *Orientia tsutsugamushi* is an obligate endosymbiont, and the five species *Symbiobacterium thermophilum*, *Thermoanaerobacter wiegelii*, *Natranaerobius thermophilus*, *Thermobacillus composti*, and *Thermosynechococcus elongatus* are thermophiles. By contrast, species with large numbers of bacterial-C-type G2Is include some that are not classified as obligate symbiotic bacteria or thermophiles. Therefore, other factors may also be involved in the increase in G2Is.

**Fig. 3.**
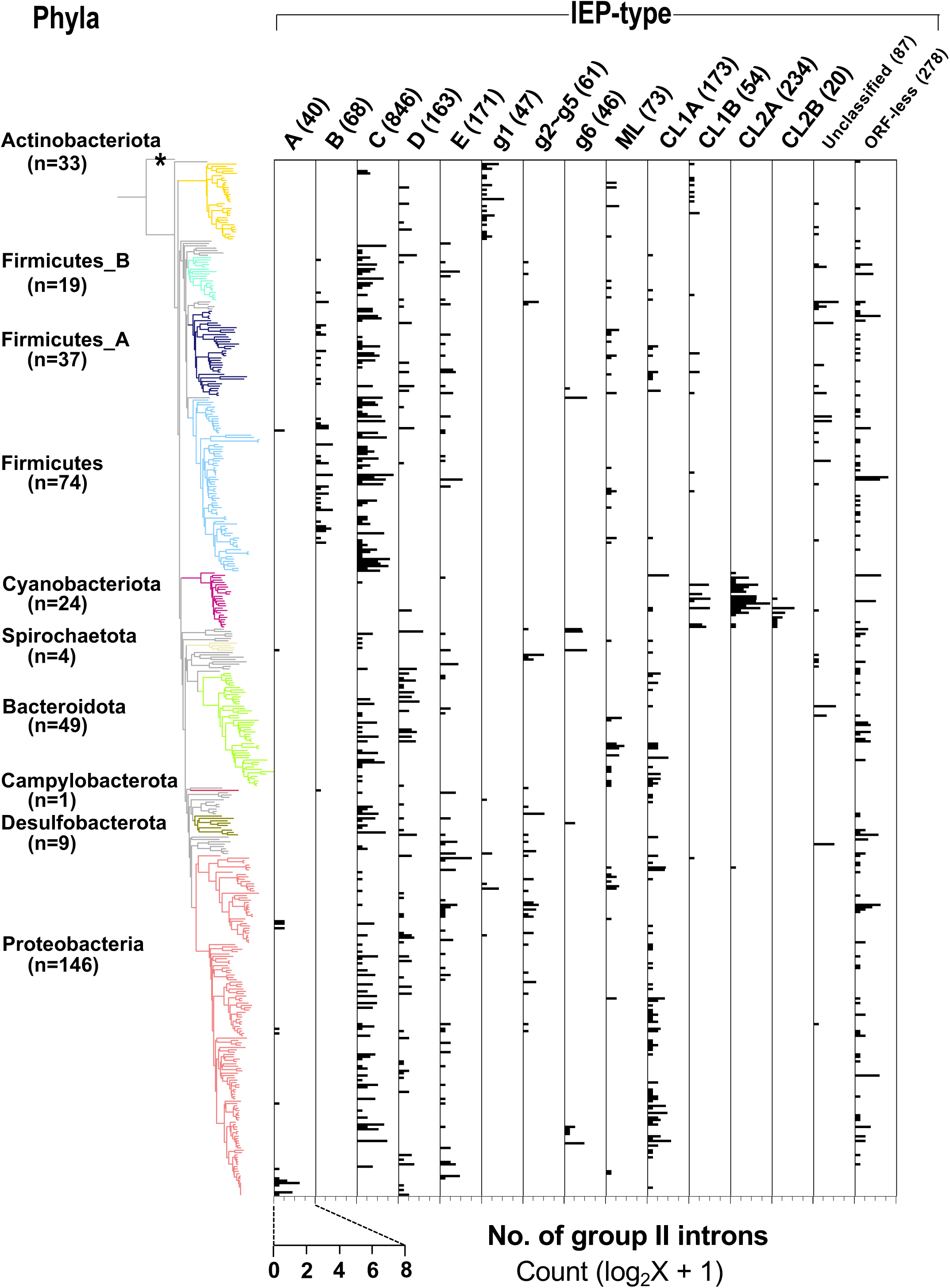
Distribution of types of IEPs in the bacterial phylogeny. A bacterial phylogenetic tree was constructed from 443 representative bacterial species whose genome contain G2I(s) (see Materials and Methods), and *Candidatus Saccharibacteria* oral taxon TM7x (RefSeq assembly accession: GCF_000803625.1) was used as the outgroup. See the legends of Figures 1 and 2 for details. The horizontal axis represents the number of G2Is for each IEP type.

Our analysis has so far focused on the genomes of representative bacterial species. Therefore, we examined the phylum Cyanobacteriota because it contains species with the largest numbers of G2Is per genome, and information on the genomes of 130 cyanobacterial species, including nonrepresentative species, is available. Supplementary Figure S10 shows the numbers of G2Is and the types of IEPs on a phylogenetic tree of 130 species from the phylum Cyanobacteriota. It shows that CL-type G2Is occur in the majority of Cyanobacteriota species with G2Is. The families with species containing ≥ 20 G2Is were Thermosynechococcaceae, Rubidibacteraceae, Phormidiaceae, and Coleofasciculaceae. It is noteworthy that the numbers of G2Is varied among the species and strains within these families, and some species had a small number of G2Is (≤ 5). Within the phylum Cyanobacteriota, members of the family Phormidiaceae have large numbers of G2Is. For example, in this family, species belonging to the genus *Arthrospira* have about 30–100 G2Is, and two of the three species belonging to the other genus also have ≥ 20 G2Is. This suggests that there is a specific mechanism that increases G2Is within the family Phormidiaceae.

### Archaeal G2Is are Concentrated in the Phylum Halobacterota

We also calculated the number of G2Is per species in archaea and mapped them onto the archaeal phylogenetic tree obtained from the Genome Taxonomy Database (Supplementary Figure S11A). In the 28 archaeal species containing G2Is, the average number ± standard deviation of G2Is was 3.0 ± 2.2, and the median was 2. Archaeal G2Is were concentrated in the phylum Halobacterota. In bacteria, some species have large numbers of G2Is, sometimes ≥ 20, but in archaea, the maximum number of G2Is per genome is 10. On the IEP phylogenetic tree shown in Figure 2A, the archaeal IEPs are distributed into four clades (bacterial-C, -D, CL1A, and CL1B), and form monophyletic groups distinct from those of the bacterial IEPs (Supplementary Figure S11B). CL1A is the commonest IEP type in archaea (Supplementary Figure S11C), and among the 19 archaeal species with G2Is, 13 have CL1A-type G2Is, and *Methanosarcina siciliae* T4/M has 10 CL1A-type G2Is. With the comprehensive analysis undertaken in this study, no significant increase in bacterial-C-type G2Is, as seen in bacteria, was detected in archaea. These results are consistent with previous studies that showed a limited increase in G2Is within archaea as compared with bacteria (Simon et al. 2008).

### G2Is with Specific IEP Types Tend to Integrate Just After Rho-independent Transcription Terminators

We have shown that bacterial-C-type G2Is are widespread in bacteria and are increasing in many species. This is because bacterial-C-type G2Is are inserted immediately after rho-independent transcription terminators during transposition (Robart et al. 2007). Therefore, their effect on the host gene(s) is limited because these G2Is do not break the coding sequences in the host genome (Robart et al. 2007). Therefore, there are very few opportunities for bacterial-C-type G2Is to be transcribed (Robart et al. 2007; Mohr et al. 2018). However, the relationship between bacterial-C-type G2Is and transcription termination sites has only been suggested in a limited number of cases. To confirm the positional relationship between the G2Is that we identified and previously predicted rho-independent transcription terminators (Mitra et al. 2011), the distances from the 5′ ends of the G2Is to the transcription termination sites were calculated for 304 bacterial species with G2Is (Figure 4). There are no data on the 5′-end consensus sequence of the bacterial-g1-type G2Is (see Materials and Methods), and the 5′-end sequence could not be determined, even when the similarity of G2Is to other IEP types was used, so bacterial-g1-type G2Is were excluded from the present study. Our results showed that, in the bacterial-C type G2Is, the distribution of the distance peaked at 0–50 bases. Thus, bacterial-C-type G2Is are dominated by sequences inserted immediately after the transcription terminator. As in the case of the bacterial-C-type G2Is, nearly half the G2Is that occurred immediately after the transcription terminator contained other common IEP types, such as bacterial-g6 and unclassified U1 and U2. In this analysis, the data available for the region of transcription termination were limited, and the data on unclassified types of IEPs referred to only U1 and U2. In G2Is inserted immediately after the transcription terminator, the ratio of U1 IEP and U2 IEP was almost the same, and there was no bias. The bacterial-C, g6, U1, and U2 types of IEP were located close to each other on the IEP phylogenetic tree (Figure 2A). In contrast, the g2–g5-type IEPs did not map directly after transcription termination regions in the bacterial genomes, even though they were surrounded by the C, g6, U1, and U2 types on the IEP phylogenetic tree. These results suggest that the regulation of G2I insertion changed during the evolution of the g2–g5-type IEPs, although the reason for this change is not clear. By contrast, other IEP-type G2Is tended to be located more than 1,000 bases from the transcription termination sites. We thought that these G2Is can enter the coding sequences of gene and cause disruption, or occur in intergenic regions, without a rho-independent transcription terminator. We next examined how the type of transcription terminator contributes to the increase in G2Is for each IEP type. There are two major types of rho-independent transcription terminators, L-shaped and I-shaped. The L-shaped transcription terminator is defined as the presence of four or more U residues in the tail region immediately after the stem–loop structure of the 3′ untranslated region (UTR) of the transcript. All of the other transcription terminators are classified as I-shaped (Mitra et al. 2011). No clear tendency was observed for any IEP type to insert preferentially into the L-shaped or the I-shaped transcription termination regions (Figure 4).

**Fig. 4.**
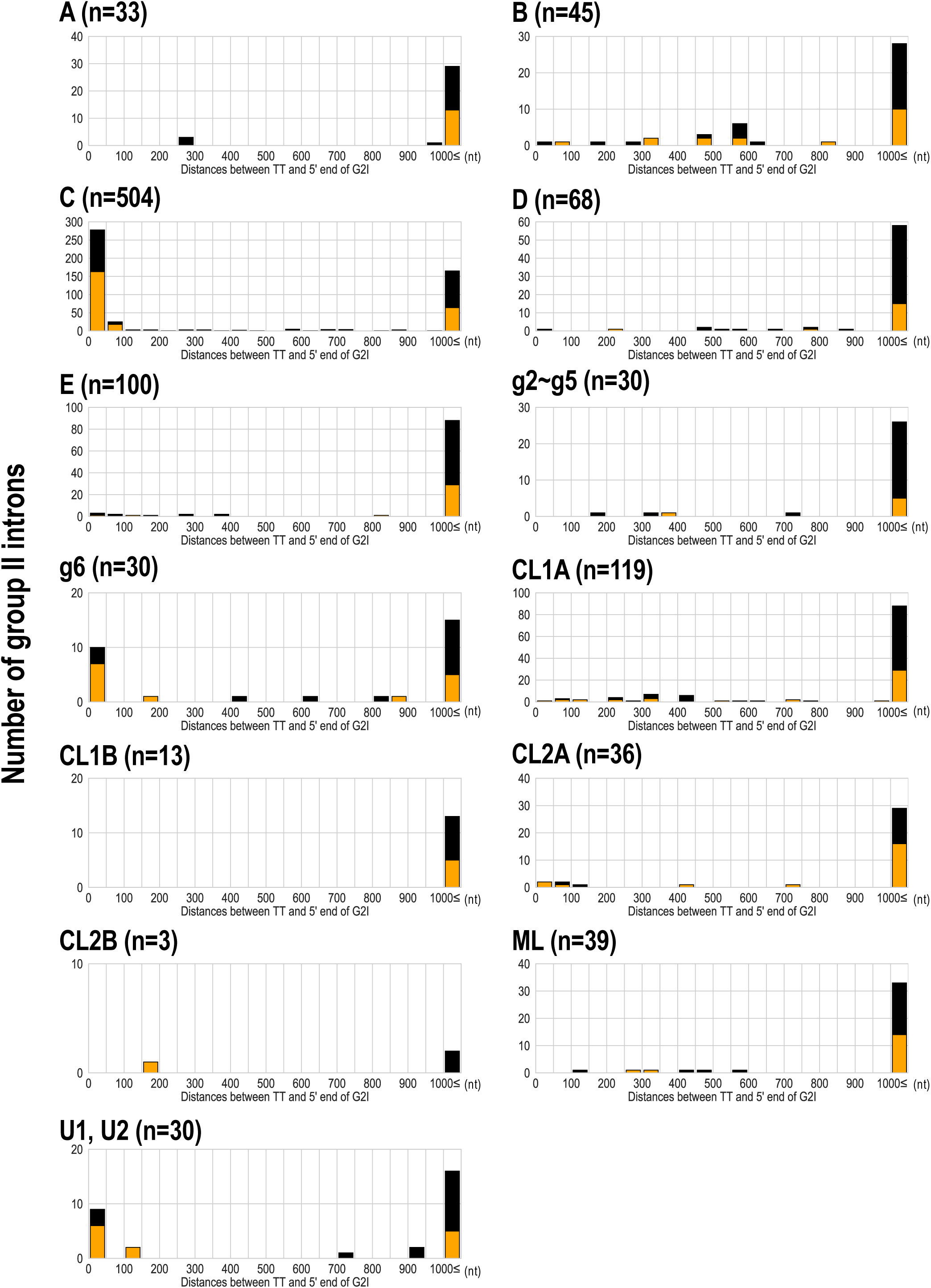
Analysis of the distance between the transcriptional terminator and the 5′ end of G2Is for each IEP type in bacteria. The distance between the transcriptional terminator and the 5′ end of the G2I was calculated, and the number of G2Is at each distance is represented as a histogram for each IEP type. L-shaped terminators are shown in orange boxes and I-shaped terminators are shown in black boxes. G2Is with bacterial-g1 IEPs are not shown because their 5′ ends were not identified in this analysis. TT: transcriptional terminator.

### The DNA Strand into Which G2Is Insert Differs for Each IEP Type

In addition to the structure of the transcription terminator, the DNA strand into which the G2I inserts is another factor that may influence the numbers of G2Is for each IEP type. It has been shown that bacterial-C-type G2Is are preferentially inserted into the template strand for lagging-strand synthesis, which exposes more single-stranded DNA regions during host DNA replication (Robart et al. 2007). Strand-specific insertion has been reported even when there is no insertion downstream from the transcription terminator in the genome. A well-studied G2I, Ll.LtrB, with an ML-type IEP has been shown to be strand-specifically inserted into replication forks in the retrotransposition pathway (Ichiyanagi et al. 2002). These strand-specific transitions are also known in many transposable elements other than G2Is (Ton-Hoang et al. 2010; Gomez et al. 2014). Therefore, to examine the possibility that each IEP type is inserted into a specific DNA strand, we investigated whether G2Is occur on the leading strand or lagging strand in genomes for which information on the origin of DNA replication (*ori*) was available. The insertion bias (IB) score was calculated as the ratio of the number of G2Is on the leading strand to the number of G2Is on the lagging strand, and the bias of the insertion strand was quantified. In this way, we found that the insertion strand differed for each IEP type: G2Is with bacterial-B, -C, g2–g5, g6, or unclassified IEP types were strongly distributed on the leading strand (IB score ≥ 5), whereas those with bacterial-D, -E, ML, or ORF-less IEP types were moderately distributed on the leading strand (IB score ≥ 2 to < 5) (Supplementary Figure S12). Although similar results were reported by Zimmerly’s laboratory (Wu 2018), we quantitatively evaluated the G2I strand bias using the IB score.

### Integration of Most G2Is is Associated with the IEP Type and GC Skew Bias in the Bacterial Genome

If the DNA strand in which G2Is occur differ for each IEP type, bacteria with a single *ori* and bacteria with multiple *ori*s may contain different IEP types. This is because either the right or left replichore is the leading strand in bacteria with a single *ori*. However, the IB score analysis in the previous section was limited to species for which the position of the *ori* could be downloaded from the database. Consequently, this analysis omitted species with a characteristic genome replication mechanism in which replication starts at various positions in the genome. Therefore, we investigated the relationship between the replication mechanism and the increase in G2Is using the generalized GC skew index (GCSI) (Arakawa et al. 2009), which reflects the effect of replication bias in bacterial genomes. Here, the GC skew is a measure of strand asymmetry in the distribution of guanines and cytosines. GCSI is 1.0 if the GC skew bias in each DNA sequence is strong and 0 if no bias is present. Most bacterial species have a GCSI of about 0.1 (Supplementary Figure S13), and species with a GCSI of ≤ 0.1 do not show any clear bias in GC skew (Arakawa et al. 2009). Information of replicon and GCSI calculated in this study is summarized in Supplementary Table S5. The GCSI values varied considerably in Firmicutes, ranging from approximately 0.1 to 0.7, many towards the upper end of this range whereas those of Actinobacteriota and Cyanobacteriota tend to have low GCSI values (approximately 0–0.2) overall (Figure 5, middle row). Previous studies have suggested that some species of Cyanobacteriota have multiple *ori*s, which may be reflected in the low GCSIs of Cyanobacteriota (Watanabe 2020). Figure 5 (right row) shows that bacterial-C-type G2Is are increased in Firmicutes members with relatively high GCSIs (0.28 on average), and CL-type G2Is are increased in Cyanobacteriota members with low GCSIs (0.03 on average). This result is supported by the fact that the bacterial-C-type G2I has strong strand selectivity, whereas the CL-type G2I has weak strand selectivity (Supplementary Figure S12). In contrast, bacterial-C-type G2Is were not observed in Firmicutes species with GCSI < 0.1 (family Mycoplasmataceae), and CL-type G2Is were not observed in Cyanobacteriota species with relatively high GCSIs (approximately 0.05–0.1; family Cyanobiaceae) (Figure 5 & Supplementary Figure S10). Therefore, we analyzed the relationship between the GCSI of species containing G2Is and the number of G2Is with each IEP type. The bacterial-B- and -C-type G2Is increased when the GCSI of the host bacterium was high (0.2 or 0.08, respectively), whereas the bacterial-g1-type and CL-type G2Is increased when the host bacterium GCSI was low (we calculated the scores to be 0.01–0.07 and 0.009–0.02, respectively) (Supplementary Figure S14). To represent these findings visually for typical bacterial examples, 20 bacterial species containing G2Is were selected. We analyzed the GC skew according to the GCSI and the insertion position of each intron in the genome (Figure 6). In species with high GCSIs (approximately 0.08–0.6) and a well-defined genomic structure, bacterial-C-type G2Is were inserted unevenly on one strand. In species with lower GCSIs (approximately 0.01–0.06), such as the genera *Arthrospira* and *Desulfobacter*, G2Is other than the bacterial-C type showed no bias in the DNA strand into which it inserted. These results suggest that the transposition of G2Is occurs in a different manner for each IEP type. It occurs in association with host DNA replication, and simple species differences do not increase the number of G2Is.

**Fig. 5.**
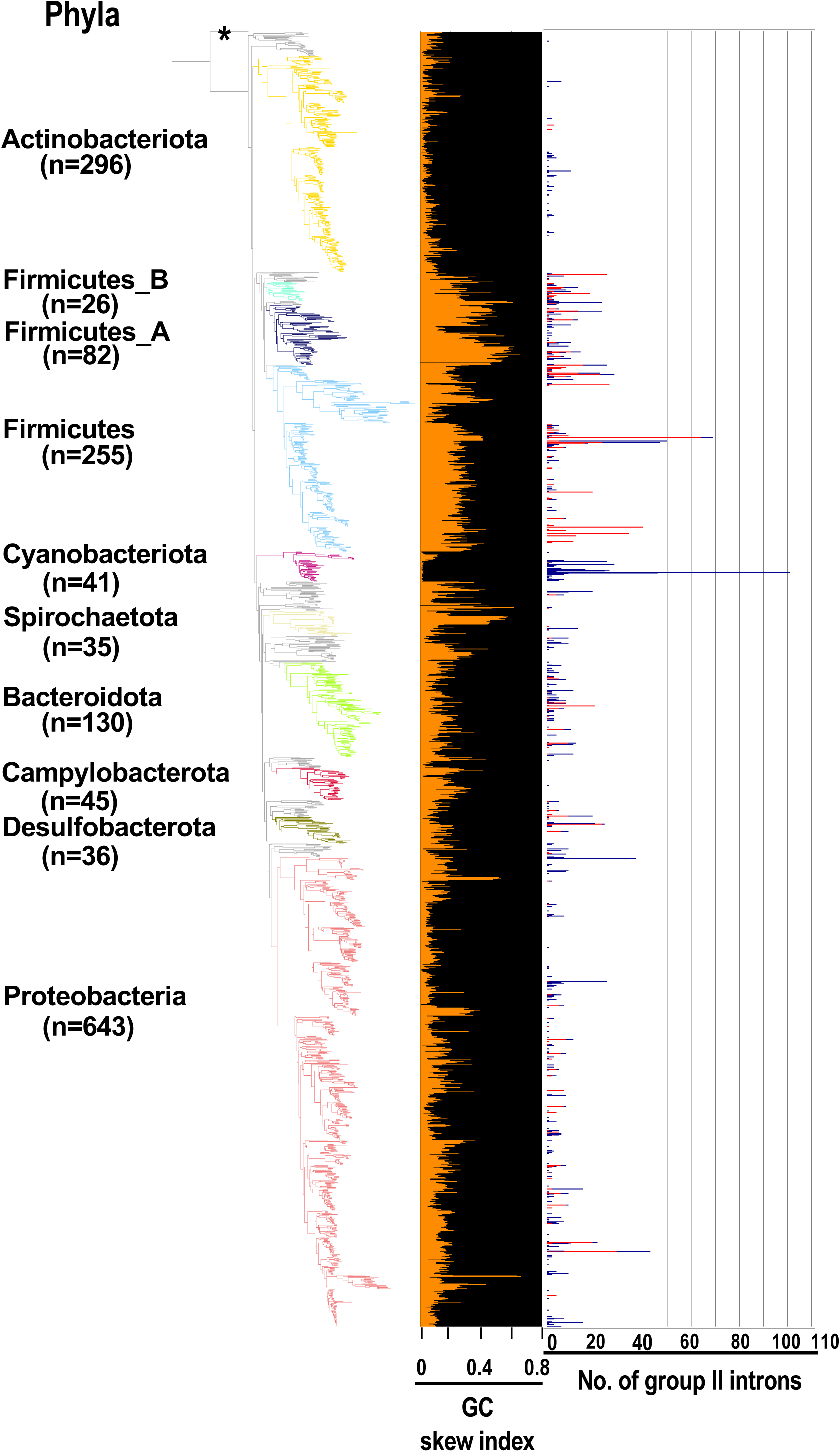
GCSI and number of G2Is in the bacterial phylogeny. The GC skew indices and numbers of G2Is in representative complete bacterial genomes (1,775 species) are shown. Bacterial phyla are shown on left and each corresponding branch on the bacterial phylogenic tree is colored. The numbers in brackets represent the number of species in each phylum. The position of the outgroup (*Candidatus Saccharibacteria* oral taxon TM7x [RefSeq assembly accession: GCF_000803625.1]) is indicated by the asterisk. The orange line in the middle panel indicates the GC skew index of the longest genome in each bacterial species. The numbers of G2Is are also shown on the right (red line: G2Is with bacterial-C type IEPs; blue line: G2Is with other IEPs).

**Fig. 6.**
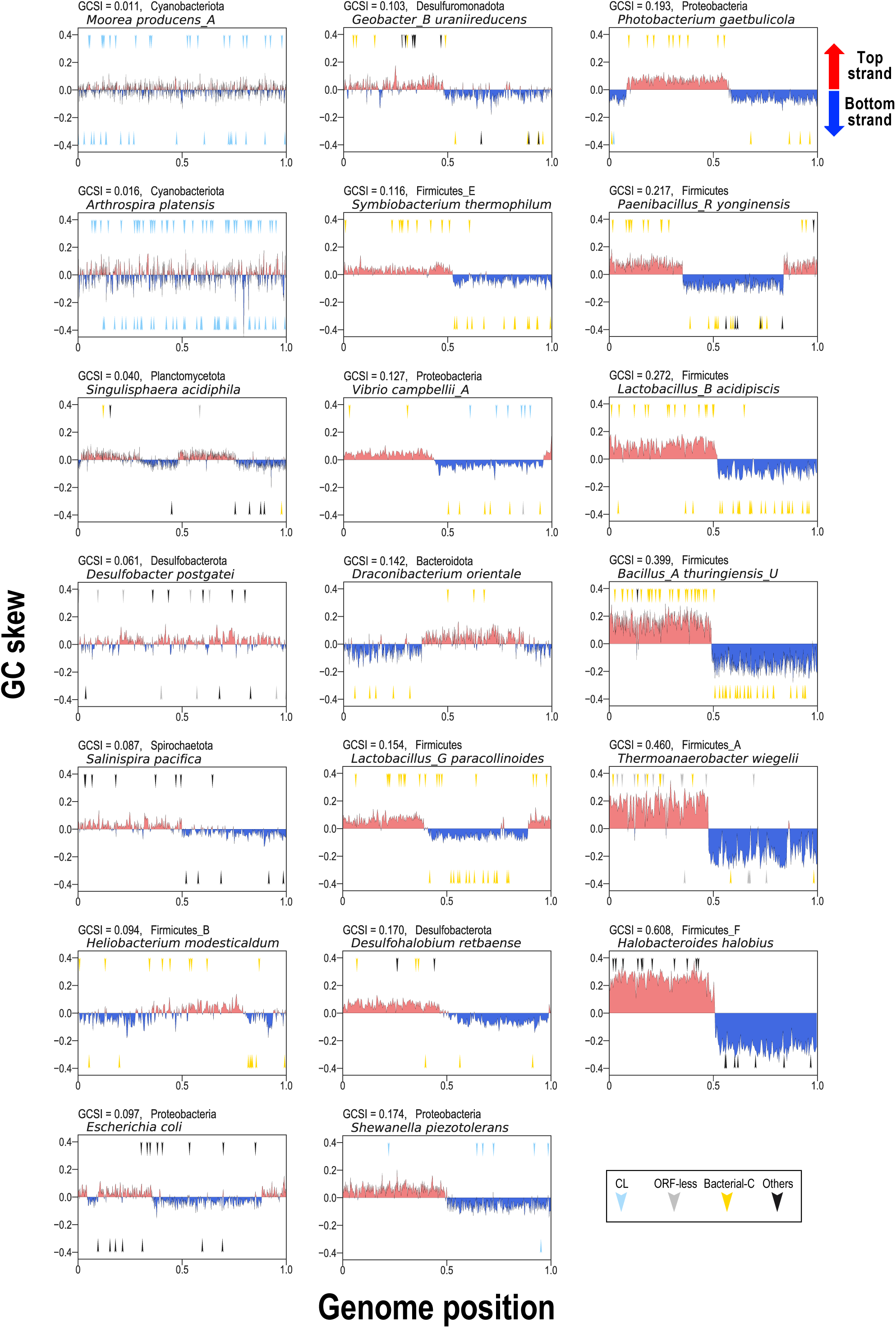
GC skew and insertion positions of G2Is in 20 representative bacterial genomes. In the boxes, the vertical axis shows the GC skew index of each genome, and the boxes are arranged in ascending order from the upper left according to the GCSI. The horizontal axis shows the relative position of each genome; the start position of the base sequence in each GenBank file is set to 0, and the end position is set to 1. The arrow in the upper half of each plot represents the insertion of G2Is into the top strand, and the arrow in the lower half represents the insertion of G2Is into the bottom strand. The colors of the arrows and the classification of G2Is are as follows: CL type (blue), ORF-less (gray), bacterial-C type (yellow), and others (black). The GCSI, bacterial phylogeny, and species name are shown in the upper left of each box. The RefSeq genome accession numbers are as follows: *Moorea producens_A*: NZ_CP017599.1; *Arthrospira platensis*: NC_016640.1; *Singulisphaera acidiphila*: NC_019892.1; *Desulfobacter postgatei*: NZ_CM001488.1; *Salinispira pacifica*: NC_023035.1; *Heliobacterium modesticaldum*: NC_010337.2; *Escherichia coli*: NC_011750.1; *Geobacter_B uraniireducens*: NC_009483.1; *Symbiobacterium thermophilum*: NC_006177.1; *Vibrio campbellii_A*: NC_009784.1; *Draconibacterium orientale*: NZ_CP007451.1; *Lactobacillus_G paracollinoides*: NZ_CP014915.1; *Desulfohalobium retbaense*: NC_013223.1; *Shewanella piezotolerans*: NC_011566.1; *Photobacterium gaetbulicola*: NZ_CP005974.1; *Paenibacillus_R yonginensis*: NZ_CP014167.1; *Lactobacillus_B acidipiscis*: NZ_LT630287.1; *Bacillus_A thuringiensis_U*: NC_022873.1; T*hermoanaerobacter wiegelii*: NC_015958.1; *Halobacteroides halobius*: NC_019978.1.

In the analysis so far, we have defined the transcription terminator, the strand into which the G2I is inserted, and the GC skew bias as factors in the prokaryotic genome that can contribute to the number of G2Is. Here, we focused on the phyla Firmicutes, which has large numbers of bacterial-C-type G2Is, and Cyanobacteriota, which has large numbers of CL-type G2Is. We first described the relationship between the rho-independent transcription terminator and the increase in bacterial-C-type G2Is in Firmicutes. Most of the bacterial-C-type G2I insertion sites are considered to occur immediately after the rho-independent transcription terminator on the leading strand (Figures 4 & 6). Compared with other phyla, Firmicutes has a higher ratio of rho-independent transcription terminators to the total number of genes, and a higher ratio of genes on the leading strand to the total number of genes (Mitra et al. 2011; Mao et al. 2012). Thus, members of Firmicutes have a higher proportion of bacterial-C-type insertion sites in their genomes than members of other phyla. However, this is only about proportions, and other phyla may have the same proportion of rho-independent transcription terminators as Firmicutes. Consequently, we speculated that in addition to the number of rho-independent transcription terminators, other factors supporting the insertion of G2Is are likely responsible for the explosive increase in bacterial-C-type G2Is in Firmicutes. Therefore, we considered the weight of the role of the rho-independent transcription terminator in Firmicutes. In *Escherichia coli* (*E. coli*) (phylum Proteobacteria), the transcription and translation processes are coupled, so RNA polymerase and the ribosome are usually linked and move at the same rate. It has been suggested that the ribosome linked to RNA polymerase inhibits the formation of RNA hairpins, resulting in the partial suppression of the majority of rho-independent transcription terminators (Wang et al. 2019). When the RNA polymerase and the ribosome are separated from each other as the movement of the ribosome slows, transcription termination by rho is promoted (Artsimovitch 2018). Therefore, rho is thought to be responsible for a wide range of transcription termination in *E. coli*. However, transcription termination in *Bacillus subtilis* (phylum Firmicutes) is less dependent on rho, and rho-independent termination is the main transcription termination mechanism (Johnson et al. 2020). This characteristic transcription termination control of *B. subtilis* is considered to be common in Firmicutes (Johnson et al. 2020). From this evidence, we infer that transcription termination by the rho-independent transcription terminator alone is predominant in Firmicutes, and when the bacterial-C-type G2I is inserted immediately after the terminator, the G2I would be rarely transcribed, suggesting that the effect of increased G2Is on the host genome is very small. It is also possible that the rho-independent transcription terminator in Firmicutes has sequence and structural features that preferentially bind to G2Is with bacterial-C-type IEPs, but further analysis is required to clarify this. By contrast, the abundance of CL-type G2Is in Cyanobacteriota may be associated with its polyploidy. Many bacteria are considered to have monoploid genomes, but some Cyanobacteriota have polyploid genomes (Griese et al. 2011). Species of Cyanobacteriota with polyploid genomes have relatively low GCSIs (Watanabe 2020). The G2I-rich Cyanobacterial species identified in our study had relatively low GCSIs and possibly polyploid genomes. For example, among the Cyanobacteriota, *Trichodesmium erythraeum* IMS101 (GCSI = 0.02) has about 100 copies of its chromosome with 24 G2Is, whereas *Synechococcus* sp. WH 8103 (GCSI = 0.085) has 1–2 chromosomes and no G2Is. In a multicopy genome, even if G2I is integrated into an essential gene on one chromosome, the gene expression from another chromosome remains intact, so the effect will be less detrimental than in haploid bacteria. The redundancy of these multicopy genomes may support the proliferation of transposable elements, such as G2I.

### Concluding Remarks

In this study, we found that the numbers of G2Is of specific IEP types have increased in specific species of bacteria, and that there are large differences in the numbers of G2Is in a wide range of bacteria, even among closely related species. The factors that determine these differences in the numbers of G2Is among closely related species have not been completely clarified. We believe that more research is required to examine the dynamics of the increases and reductions in G2Is, mainly in bacteria such as Firmicutes and Cyanobacteriota, using population genomic methods (Siguier et al. 2014; Consuegra et al. 2021). Further genomic sequencing of related species, including the analysis of ploidy, and clarification of the molecular relationship between ploidy and GC skew, will be essential, especially in Cyanobacteriota. The comprehensive analysis of IEPs in this study revealed the distribution of noncanonical IEPs. Because G2Is are thought to be inessential for the host, it seems inevitable that mutations will occur in the G2I regions of genomes, but understanding the mechanisms that act on the evolutionary coexistence of G2I and host genes awaits further research.

## Materials and Methods

### Datasets

We examined 14,506 bacterial and 296 archaeal species in this study. The genomic sequences and their annotations were downloaded in GenBank format from the NCBI RefSeq database (O’Leary et al. 2016) in March 2019. These genomic data met both of the following conditions: (i) assembly level; complete genome or chromosome; and (ii) version status; latest. We used the Biopython module (version 1.73) (Cock et al. 2009) to read the GenBank annotation data and to calculate the GC skew. Prediction data for bacterial replication origins were obtained from the DoriC database (version 10.0) (Luo and Gao 2019). Information on species used for each figure is summarized in Supplementary Table S6.

### Construction of Phylogenetic Trees for Prokaryotic Species

The phylogenetic trees of 27,372 bacterial and 1,569 archaeal species were downloaded from the Genome Taxonomy Database (version 86.2) (Parks et al. 2018) and the trees were pruned using ETE Toolkit (version 3.1.1) (Huerta-Cepas et al. 2016) to focus on specific groups of species in each analysis. The phylogenetic trees were visualized with iTOL (version 5) (Letunic and Bork 2021). *Candidatus Saccharibacteria* oral taxon TM7x, a member of the CPR bacteria (RefSeq assembly accession: GCF_000803625.1), was used as the outgroup. We constructed six phylogenetic trees for: (i) representative bacteria (1,775 species in Figure 1, Figure 5 and Supplementary Figure S4); (ii) representative bacteria containing G2I gene(s) (444 species in Figure 3); (iii) representative bacteria containing ≥ 20 G2I genes (26 species in Supplementary Figure S9) and all cyanobacterial species in the dataset (130 species in Supplementary Figure S10); (v) all archaeal species in the dataset (222 species in Supplementary Figure S11A); and (vi) all archaeal species containing G2I gene(s) in the dataset (19 species in Supplementary Figure S11C). The representative species are classified as either “representative genome” or “reference genome” in the RefSeq database (March 2019).

### Comprehensive Extraction of G2Is from Genomic Data

The pipeline used to extract G2Is from the genomic data is shown in Supplementary Figure S1A, and comprised the following steps. Step 1 (extracting three groups of domains of G2I):

i. the NCBI BLAST+ (version 2.4.0) tblastn command line tool (Camacho et al. 2009) was used to search for the RT domain of the IEP. As query sequences, we used amino acid sequences of 425 RT domains in G2I IEPs (Toro and Nisa-Martinez 2014). We set the e-value threshold to 1*e*−10 and extracted the sequences with query coverage > 40%. The query sequence with the highest bit score was then selected and the IEP type of this query sequence was defined as the IEP type of the subject sequence.
ii. To search for RNA domains V and VI of the G2I, the Infernal (version 1.1.2) program cmsearch was used (bit score threshold: > 24, -nohmm option) (Nawrocki and Eddy 2013). Here, we used the RNA secondary structure model Intron_gpII (ID: RF00029) registered in the Rfam database (http://rfam.xfam.org/) (Kalvari et al. 2018).
iii. To detect ORF-less-type G2Is, we also searched for the RNA domains I–IV of the G2Is with both the BLAST+ (version 2.4.0) blastn command line tool (*e*-value threshold: 1*e*−10) and the Infernal (version 1.1.2) program cmsearch (*e*-value threshold: 1*e*−10, -rfam option). For the blastn search, 337 G2I sequences in prokaryotes registered in the Database for Bacterial Group II Introns (http://webapps2.ucalgary.ca/~groupii/index.html#) (Candales et al. 2012) were manually selected as query sequences, and the sequences with query coverage > 60% were extracted. For the cmsearch search, we used RNA secondary structure models (IDs: RF01998, RF01999, RF02001, RF02003, RF02004, RF02005, and RF02012) registered in the Rfam database.

The following two steps were used to judge whether each detected domain was included in a single G2I. Step 2: When the distance between the regions with similarity to RT and domain V was ≤ 1,300 bases (Supplementary Figure S1B), we considered that the RT domain and domain V were in the same G2I. Step 3: When the distance between either domain I, II, III, or IV and domain V was ≤ 1,300 bases (Supplementary Figure S1C), we considered that either domain I, II, III, or IV and domain V were in the same G2I. The resulting G2I dataset was checked manually and corrected where necessary (Supplementary Table S1B, Supplementary Figure S11).

### Prediction of IEP Sequences

From our preliminary survey, it was apparent that there are noncanonical IEP sequences, such as those whose ORFs are interrupted by stop codons (Supplementary Figure S3). Therefore, in this study, the entire IEPs (or the partial sequences of IEPs) were extracted from the peripheral sequences of the identified RT domains and classified as canonical IEP or noncanonical IEP sequences. After 12,217 RT domain sequences were extracted in the previous section (“Comprehensive Extraction of G2Is from Genomic Data”), we obtained the nucleotide sequences corresponding to the 1,200 bases upstream from the RT domain to the 200 bases downstream from domain VI in each genomic sequence (Supplementary Figure S1A, Step 1). We then searched for the nucleotide sequences corresponding to IEP amino acid sequences around these RT domains with the tblastn command (Camacho et al. 2009). Each subject sequence with the highest bit score was selected, and a total of 12,217 IEP amino acid sequences were obtained. Here, the amino acid sequences of each IEP of the 318 “Eubacterial” G2Is registered in Zbase was used as queries (Candales et al. 2012). We selected 1,086 sequences in which a stop codon appeared other than at the end of the subject sequence, and called these sequences “interrupted sequences”. To reduce the number from these 1,086 sequences before the construction of a phylogenetic tree, clustering was performed with CD-HIT (version 4.8.1) (Fu et al. 2012) with a threshold of 85% sequence similarity, and 360 clusters were extracted. By selecting one representative sequence from each cluster, a representative sample of 360 “interrupted sequences” was obtained.

For the 11,130 IEP sequences that were not “interrupted sequences”, the ORFs and the conserved domains in each IEP were predicted, and if the conserved functional domain was missing, it was considered a noncanonical IEP. That is, the ORF was predicted with ORFfinder (https://www.ncbi.nlm.nih.gov/orffinder/) around each RT domain in these 11,130 nucleotide sequences. This process was the source of the 11,130 IEP sequences. Because there are many frames in an ORF that differ from that of the IEP, the IEP sequences in these ORFs were selected with the blastp command, and 11,130 IEP ORFs were obtained. Next, because the number of “interrupted sequences” was large, it was difficult to create a phylogenetic tree and analyze the missing domains, so clustering was performed again using CD-HIT (version 4.8.1) with a threshold of 85% sequence similarity. This yielded 1,617 clusters. Using MEME Suite (Bailey et al. 2009), a total of 15 conserved sequence domains with lengths of 10–50 amino acids were set for each IEP type, and those missing about 5 or more domains were manually selected as IEPs with deleted domains. Consequently, 210 ORFs with large domain deletions were obtained, and these ORFs were called “short ORFs”. Finally, 570 ORFs (the sum of “interrupted sequences” and “short ORFs”) were designated “noncanonical IEPs”, and the remaining 1,407 ORFs were designated “canonical IEPs”.

### Construction of Phylogenetic Tree of Intron-encoded Proteins

The dataset used contained 1,977 IEP sequences, including 1,407 sequences of canonical IEPs and 570 sequences of noncanonical IEPs (see “Prediction of IEP Sequences”). If the RT domain sequence of a noncanonical IEP was interrupted by a stop codon, the relevant stop codon was excluded. To construct a phylogenetic tree of the IEPs, MAFFT E-INS-i (version 7.310) (Katoh and Standley 2014) was used to prepare a multiple alignment of the 1,977 IEP sequences, which was trimmed with trimAl (version 1.2, gappyout option) (Capella-Gutierrez et al. 2009). The phylogenetic tree was constructed with the maximum likelihood method with RAxML (version 8.2.10) (Stamatakis 2014) using the LG + Γ model and 100 bootstrap replicates (raxmlHPC-PTHREADS-SSE3 -f a -N 100 -m PROTGAMMALG). The phylogenetic tree was visualized with iTOL (version 5).

### Calculation of Distances from G2I Insertion Sites to Transcription Termination Sites

To determine which IEP-type G2Is are inserted immediately after the rho-independent transcription terminator, the distances between the rho-independent transcription terminators and the 5′ ends of the G2Is in the bacterial genomes were calculated. First, prediction data for rho-independent transcription terminators that included information on the type of terminator (L-shaped or I-shaped) were downloaded from the WebGeSTer DB (last updated: June 06, 2012) (Mitra et al. 2011), and used as “the best or strongest candidate terminators.” Because the 5′ end of a G2I is expected to be located 5′ upstream from the start codon of an IEP and within about 300–1,100 bases of it (Supplementary Figure S1), we searched for the 5′ end sequence within the region within 1,200 bases upstream from the IEP. Because files of the 5′ end consensus sequence of G2Is have already been published for each of the nine IEP types (bacterial-A, -B, -C, -D, -E, -F [g2–g5], ML, CL1, and CL2) (Waldern et al. 2020), we searched for regions with similarity to these consensus sequences within the 1,200 bases upstream from the IEPs. The positions of the 5′ ends were then mapped onto the genome of each bacterium with the following method. Using the hmmbuild and hmmpress commands of HMMER (version 3.1b2), a hidden Markov model (HMM) profile was created for each IEP type from multiple-alignment-containing files of consensus sequences (Mistry et al. 2013; Wheeler and Eddy 2013). The nhmmscan command was then used to identify the 5′ end. For this, the default values were used for the parameters, and only hits with a bit score of ≥ 10 were selected. If there were multiple 5′ end candidates, the position with the highest bit score was selected. Finally, the distance from the 5′ end of the G2I to the 3′ end of the rho-independent transcription terminator was calculated.

### GC Skew Analysis

The GC skew of each genome was calculated as (G − C)/(G + C), where G and C represent the numbers of guanines and cytosines, respectively, in windows of 10,000 bp, using the Biopython module (version 1.73) (Cock et al. 2009). The GC skew index, which represents the strength of the GC skew, was calculated using the G-language Genome Analysis Environment (version 1.9.1) (Arakawa et al. 2003; Arakawa et al. 2009). To construct Figure 6, we manually selected 20 representative bacterial chromosomes containing relatively many G2Is to clarify whether differences in the GC skew indices of genomes affect the genomic region into which G2Is are inserted.

### Single Regression Analysis

To calculate the correlation coefficient between the genome size and the number of G2Is, a single regression analysis was performed using scikit-learn 0.23.1 (*sklearn.linear_model.LinearRegression*) (Pedregosa et al. 2011).

## Acknowledgments

This work was supported in part by research funds from the Yamagata Prefectural Government and Tsuruoka City, Japan. The funding bodies played no role in the study design, the data collection or analysis, the decision to publish, or the preparation of the manuscript. The authors thank all the members of the RNA Group at the Institute for Advanced Biosciences of Keio University, Japan, for their insightful discussions.

## Data Availability Statements

The data underlying this article are available in the article and in its online supplementary material.

