## Supplementary material for "Distinct Expansion of Group II Introns Depends on the Type of Intron-encoded Protein and Genomic Signatures in Prokaryotes": Suppl_Info_I

### Supplementary Information I

**Supplementary Table S1**

**Supplementary Fig. S1 - Supplementary Fig. S8**

#### Distinct Expansion of Group II Introns Depends on the Type of Intron-encoded Protein and Genomic Signatures in Prokaryotes

Masahiro C. Miura,[1,2] Shohei Nagata,[1] Satoshi Tamaki,[1]
Masaru Tomita,[1,2,3] and Akio Kanai[1,2,3*]

[1]Institute for Advanced Biosciences, Keio University, Tsuruoka, Japan
[2]Systems Biology Program, Graduate School of Media and Governance, Keio University, Fujisawa, Japan
[3]Faculty of Environment and Information Studies, Keio University, Fujisawa, Japan

*Corresponding Author

Akio Kanai, PhD
Institute for Advanced Biosciences, Keio University
Tsuruoka, Yamagata, Japan


**List of abbreviations:**

| | |
|---|---|
| CPR | Candidate Phyla Radiation |
| E1 | exon 1 |
| E2 | exon 2 |
| FN | false negative |
| FP | false positive |
| GCSI | GC skew index |
| G2I | group II intron |
| IB | insertion bias |
| IEP | intron-encoded protein |
| Mbp | megabase pair |
| NCBI | National Center for Biotechnology Information |
| ORF | open reading frame |
| *ori* | origin of DNA replication |
| RefSeq | reference sequence |
| RT domain | reverse transcriptase domain |
| TP | true positive |

**Supplementary Table S1. Summary of the performance evaluation of our bioinformatic pipeline used to extract G2Is from prokaryotic genomes.**

| Domain | NCBI assemnbly ID | Organisms | No. of G2Is in reference | No. of G2Is detected by our program | TP | FP | FN | Sensitivity | Precision | Reference |
|---|---|---|---|---|---|---|---|---|---|---|
| Bacteria | GCF_000009045 | *Bacillus subtilis* 168 | 0 | 0 | 0 | 0 | 0 | - | - | - |
| Bacteria | GCF_000068525 | *Chlamydia trachomatis* L2b/UCH-1/proctitis | 0 | 0 | 0 | 0 | 0 | - | - | - |
| Bacteria | GCF_000007785 | *Enterococcus faecalis* V583 | 1 | 2 | 1 | 1 | 0 | 1.00 | 0.50 | Candales, *et al.* 2012 |
| Bacteria | GCF_000005845 | *Escherichia coli* K-12 strain MG1655 | 0 | 0 | 0 | 0 | 0 | - | - | - |
| Bacteria | GCF_000008525 | *Helicobacter pylori* 26695 | 0 | 0 | 0 | 0 | 0 | - | - | - |
| Bacteria | GCF_000196035 | *Listeria monocytogenes* EGD | 0 | 0 | 0 | 0 | 0 | - | - | - |
| Bacteria | GCF_000009445 | *Mycobacterium bovis* BCG Pasteur | 0 | 0 | 0 | 0 | 0 | - | - | - |
| Bacteria | GCF_000006765 | *Pseudomonas aeruginosa* PAO1 | 0 | 0 | 0 | 0 | 0 | - | - | - |
| Bacteria | GCF_000007545 | *Salmonella enterica* subsp. enterica serovar Typhi Ty2 | 0 | 0 | 0 | 0 | 0 | - | - | - |
| Bacteria | GCF_000009645 | *Staphylococcus aureus* N315 | 0 | 0 | 0 | 0 | 0 | - | - | - |
| Bacteria | GCF_000011765 | *Streptococcus pyogenes* MGAS5005 | 0 | 0 | 0 | 0 | 0 | - | - | - |
| Bacteria | GCF_000006745 | *Vibrio cholerae* El Tor | 0 | 0 | 0 | 0 | 0 | - | - | - |
| Bacteria | GCF_000008005 | *Bacillus cereus* ATCC 10987 | 7 | 7 | 7 | 0 | 0 | 1.00 | 1.00 | Tourasse, *et al.* 2008 |
| Bacteria | GCF_000008765 | *Clostridium acetobutylicum* ATCC 824 | 1 | 1 | 1 | 0 | 0 | 1.00 | 1.00 | Candales, *et al.* 2012 |
| Bacteria | GCF_000006965 | *Sinorhizobium meliloti* 1021 | 6 | 6 | 6 | 0 | 0 | 1.00 | 1.00 | Toro, *et al.* 2003 |
| Bacteria | GCF_000011345 | *Thermosynechococcus elongatus* BP-1 | 28 | 25 | 25 | 0 | 3 | 0.89 | 1.00 | Nakamura, *et al.* 2002 |
| Bacteria | GCF_000013645 | *Paraburkholderia xenovorans* LB400 | 3 | 4 | 3 | 1 | 0 | 1.00 | 0.75 | Candales, *et al.* 2012 |
| Bacteria | GCF_000380335 | *Azotobacter vinelandii* CA | 7 | 9 | 7 | 2 | 0 | 1.00 | 0.78 | Candales, *et al.* 2012 |
| Bacteria | GCF_000073005 | *Wolbachia* endosymbiont of Culex quinquefasciatus Pel strain wPip | 6 | 5 | 5 | 0 | 1 | 0.83 | 1.00 | Leclercq, *et al.* 2011 |
| Bacteria | GCF_000014625 | *Pseudomonas aeruginosa* UCBPP-PA14 | 1 | 1 | 1 | 0 | 0 | 1.00 | 1.00 | Candales, *et al.* 2012 |
| Bacteria | GCF_000014545 | *Lactococcus lactis* subsp. cremoris SK11 | 2 | 2 | 2 | 0 | 0 | 1.00 | 1.00 | Candales, *et al.* 2012 |
| Archaea | GCF_000013725 | *Methanococcoides burtonii* DSM 6242 | 4 | 8 | 4 | 4 | 0 | 1.00 | 0.50 | Candales, *et al.* 2012 |
| Archaea | GCF_000006805 | *Halobacterium salinarum* NRC-1 | 0 | 0 | 0 | 0 | 0 | - | - | - |
| Archaea | GCF_000970205 | *Methanosarcina mazei* S-6 | 1 | 2 | 1 | 1 | 0 | 1.00 | 0.50 | Candales, *et al.* 2012 |
| Archaea | GCF_000025685 | *Haloferax volcanii* DS2 | 0 | 0 | 0 | 0 | 0 | - | - | - |
| Archaea | GCF_000009965 | *Thermococcus kodakarensis* KOD1 | 0 | 0 | 0 | 0 | 0 | - | - | - |
| Archaea | GCF_000007305 | *Pyrococcus furiosus* DSM 3638 | 0 | 0 | 0 | 0 | 0 | - | - | - |
| Archaea | GCF_000017225 | *Methanococcus maripaludis* C7 | 0 | 0 | 0 | 0 | 0 | - | - | - |
| Archaea | GCF_000012285 | *Sulfolobus acidocaldarius* DSM 639 | 0 | 0 | 0 | 0 | 0 | - | - | - |
| Archaea | GCF_900079115 | *Saccharolobus solfataricus* P1 | 0 | 0 | 0 | 0 | 0 | - | - | - |

TP, true positive; FP, false positive; FN, false negative.

**Supplementary Table S2. Summary of G2Is detected in this study.**

**Supplementary Table S3. Summary of prokaryotic genomes used in this study.**

**Supplementary Table S4. Number of G2Is by IEP type in each species.**

**Supplementary Table S5. Information of replicon and GC skew index.**

**Supplementary Table S6. Summary of species used for each figure.**

These Tables are supplied as a separate Excel file.

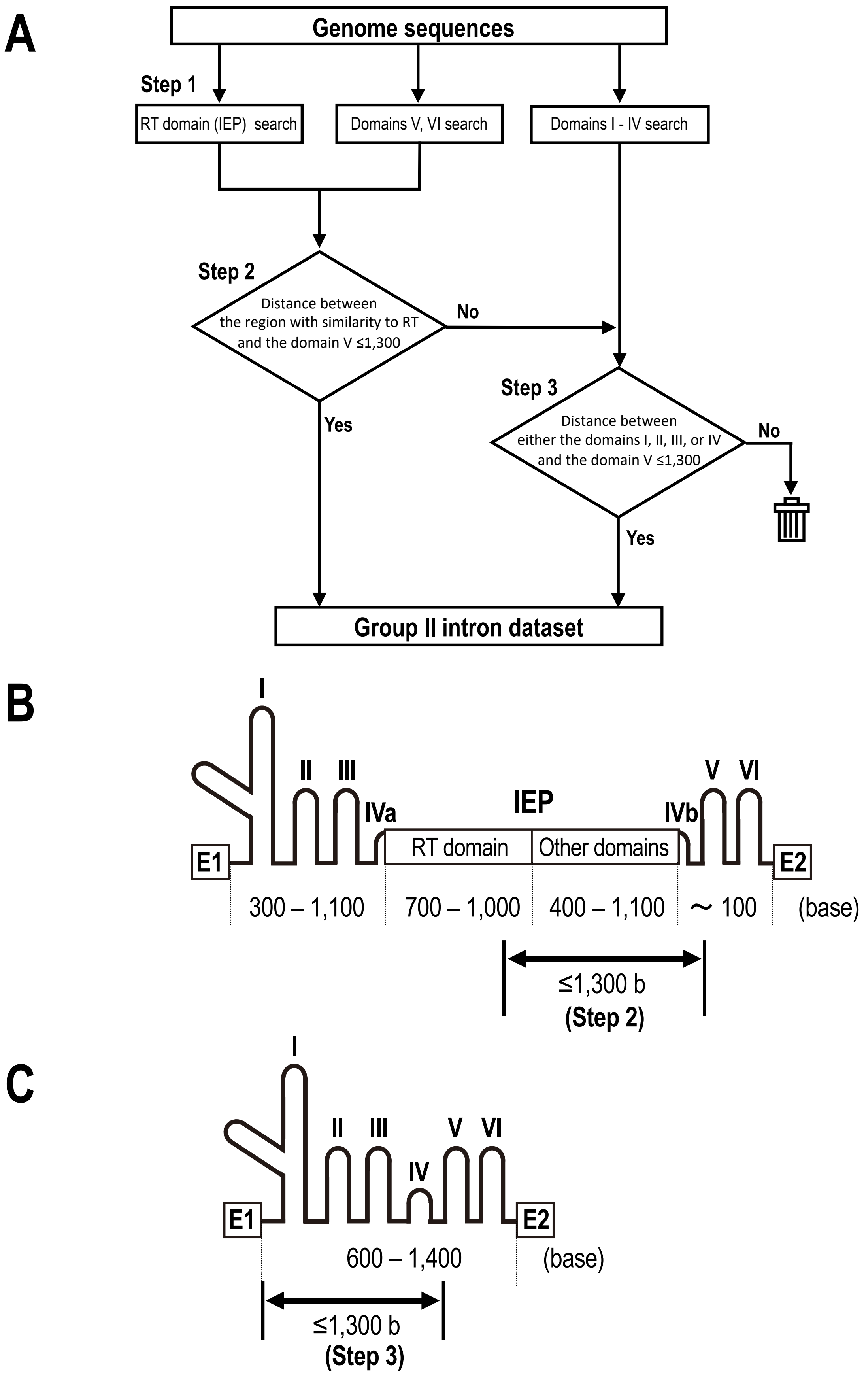


**Supplementary Fig. S1.** Bioinformatic pipeline used to extract G2Is from genomes. (A) Basic pipeline used to extract G2I datasets from genomes. (B) Graphical explanation of step 2 in the pipeline. (C) Graphical explanation of step 3 in the pipeline. IEP, intron-encoded protein; RT domain, reverse transcriptase domain; E1, exon 1; E2, exon 2. See Materials and Methods for details.

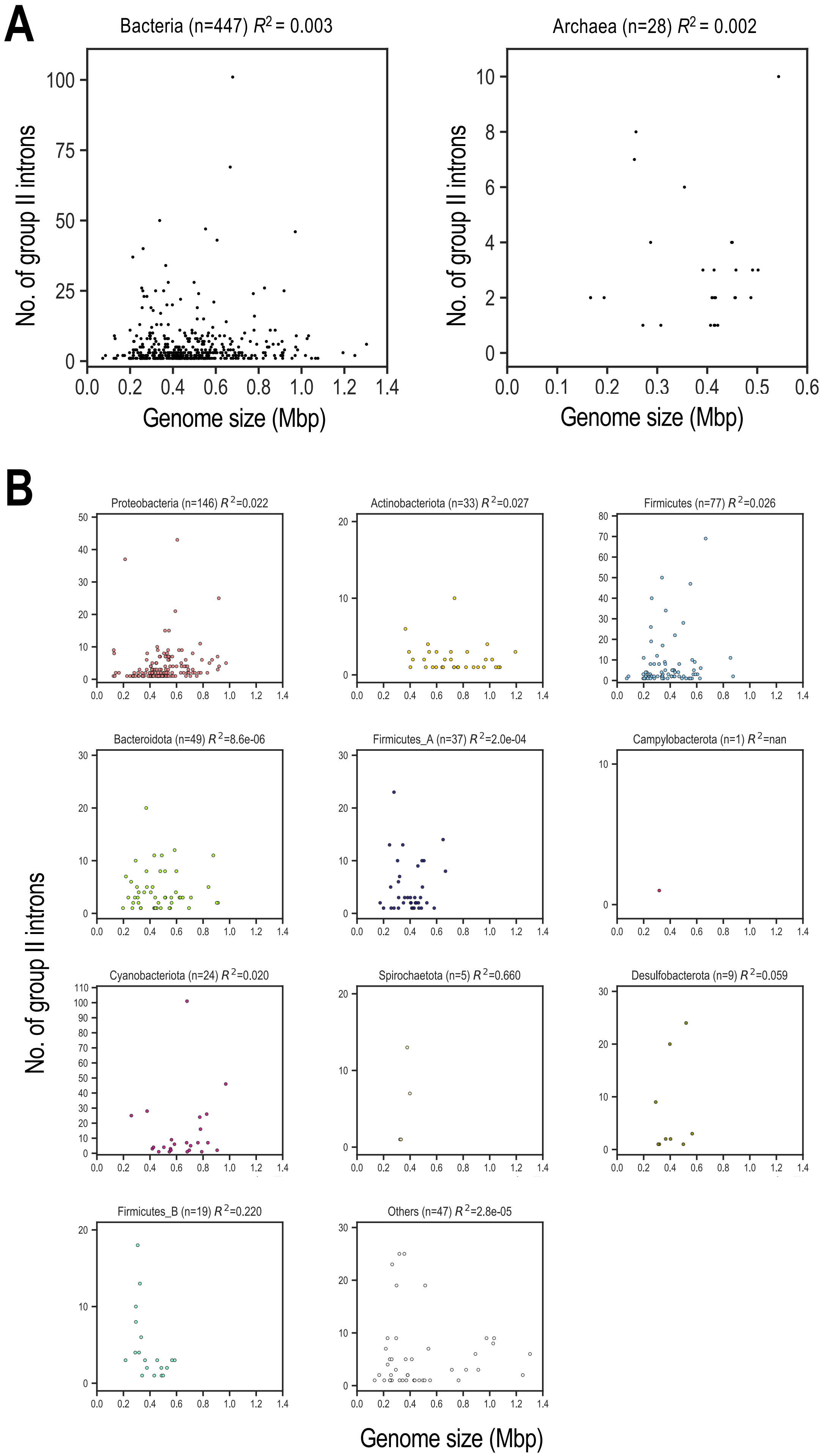


**Supplementary Fig. S2.** No correlation between numbers of G2Is in genomes and genome size. (A) Each dot indicates individual species in bacteria or archaea. (B) Each dot represents individual species in each bacterial phylum. The numbers in brackets indicate the number of species containing G2I(s). $R^2$ is the correlation coefficient in a single regression analysis.

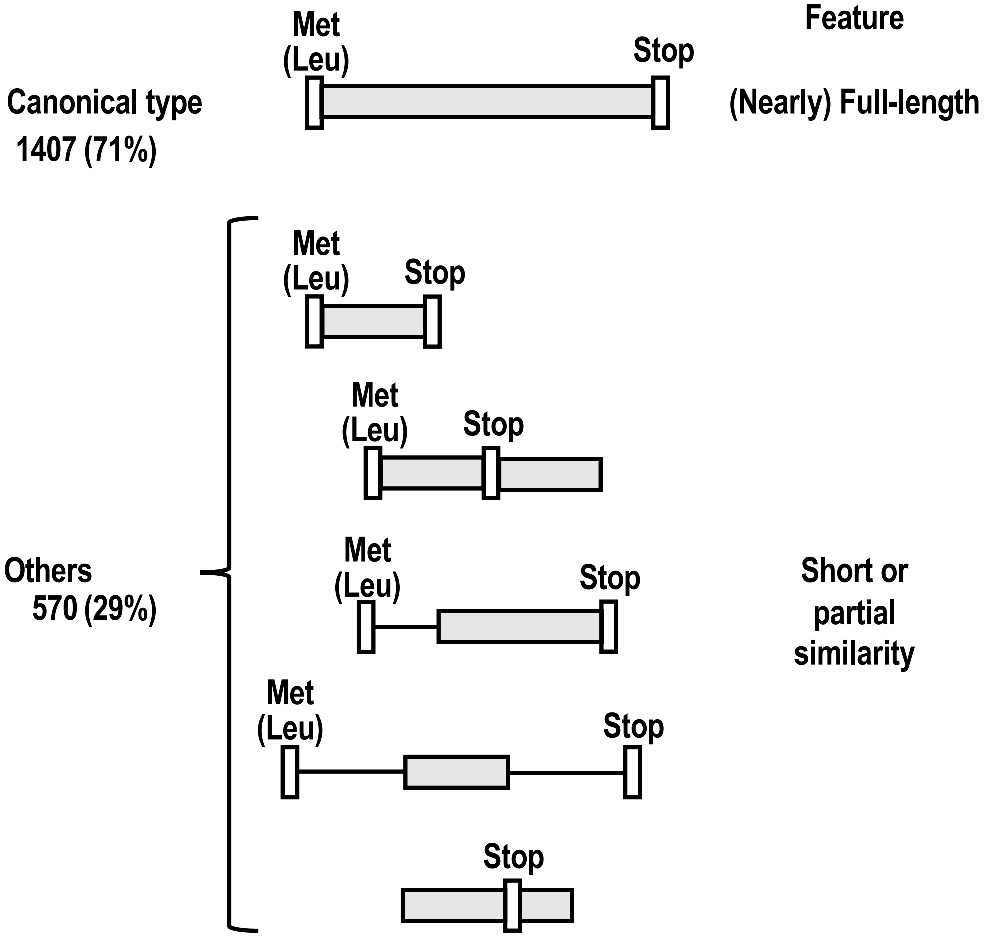


**Supplementary Fig. S3.** Schematic representation of canonical IEPs and noncanonical IEPs (others). Gray boxes indicate regions with similarity to the queried IEP and black lines indicate regions with no similarity. White boxes indicate the start (either Met or Leu) and stop codons. The numbers in the left column indicate the numbers of canonical IEPs or noncanonical IEPs among 1,977 representative IEP sequences, with percentages in brackets. For details, see Materials and Methods.

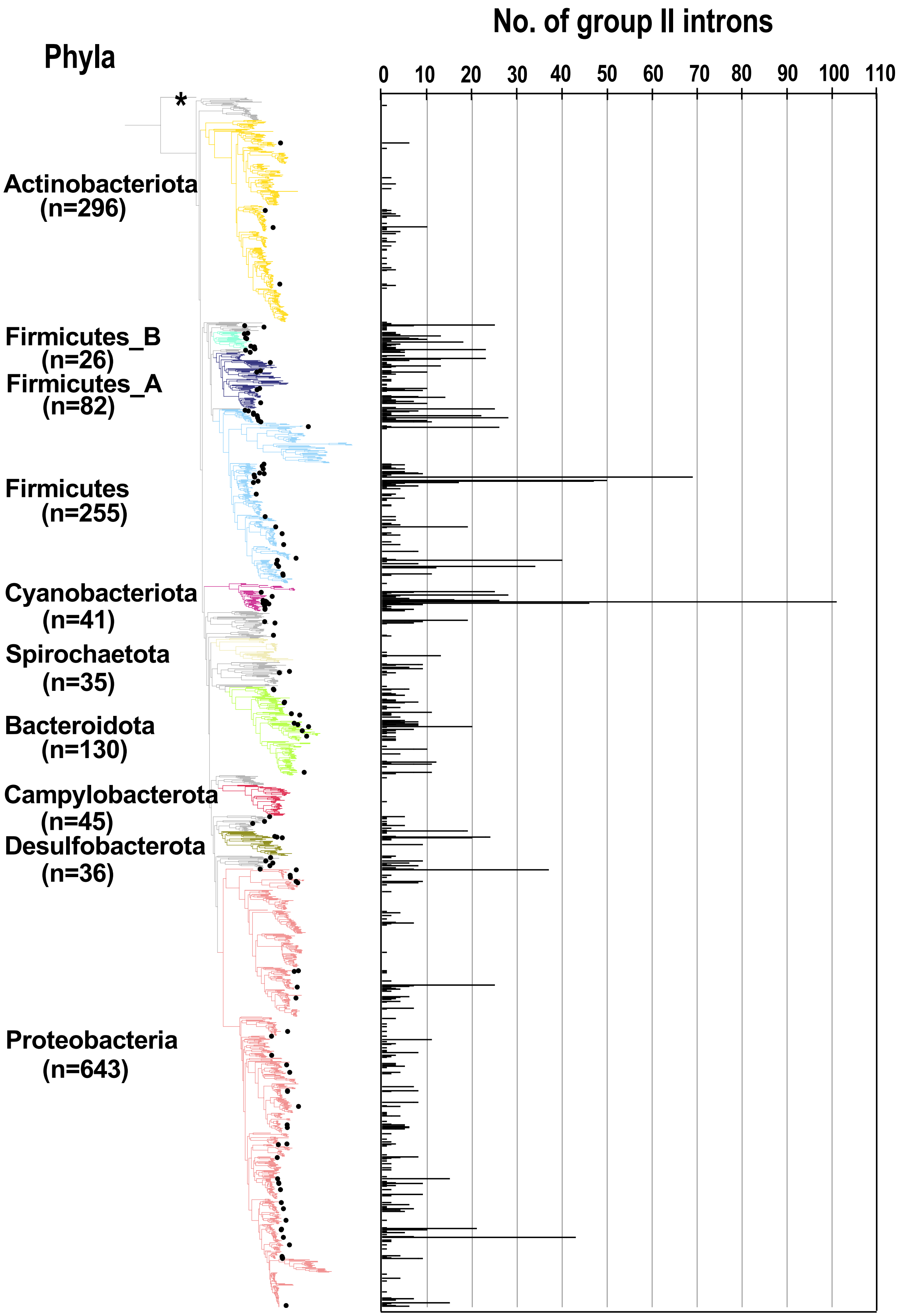


**Supplementary Fig. S4.** Distribution of bacterial species containing G2Is with noncanonical IEPs. As shown in Figure 1, numbers of G2Is in representative complete bacterial genomes (1,775 species) are shown. Dots represent the positions of species containing G2I(s) with noncanonical IEP(s). See Supplementary Figure S3 and Figure 1 legends for details.

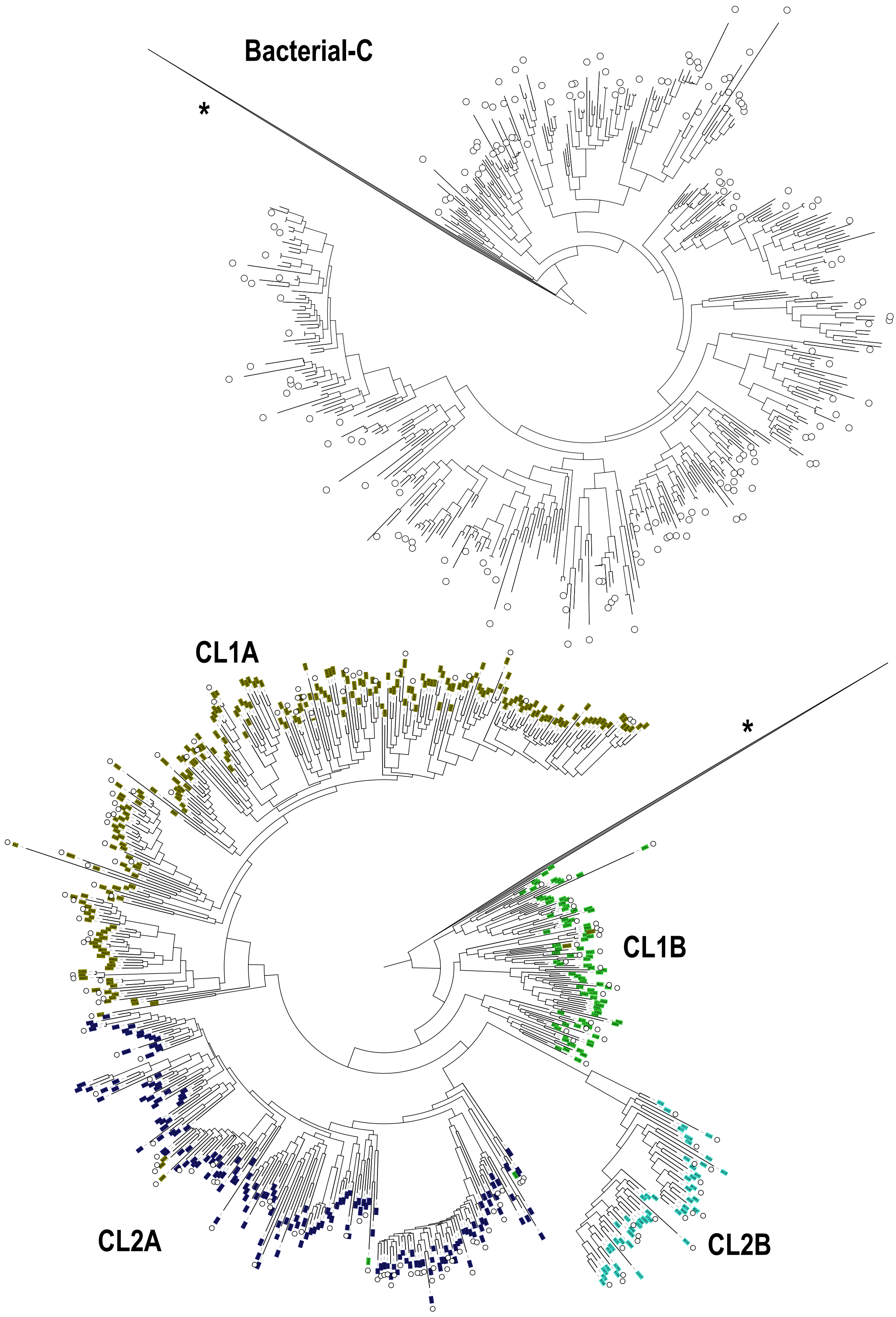


**Supplementary Fig. S5.** Distribution of noncanonical IEPs on the phylogenetic tree of the bacterial-C-type or CL-type IEPs. From the phylogenetic tree of IEPs in Figure 2A, only the clades corresponding to the bacterial-C type (upper figure) or CL type (lower figure) are shown. In the CL IEPs, the correspondence between color and each IEP type is as follows: CL1A (olive), CL1B (green), CL2A (blue), and CL2B (turquoise). White circles indicate positions of noncanonical IEPs. * To mainly display either the bacterial-C or CL phylogenetic tree, other branches are collapsed into a single clade.

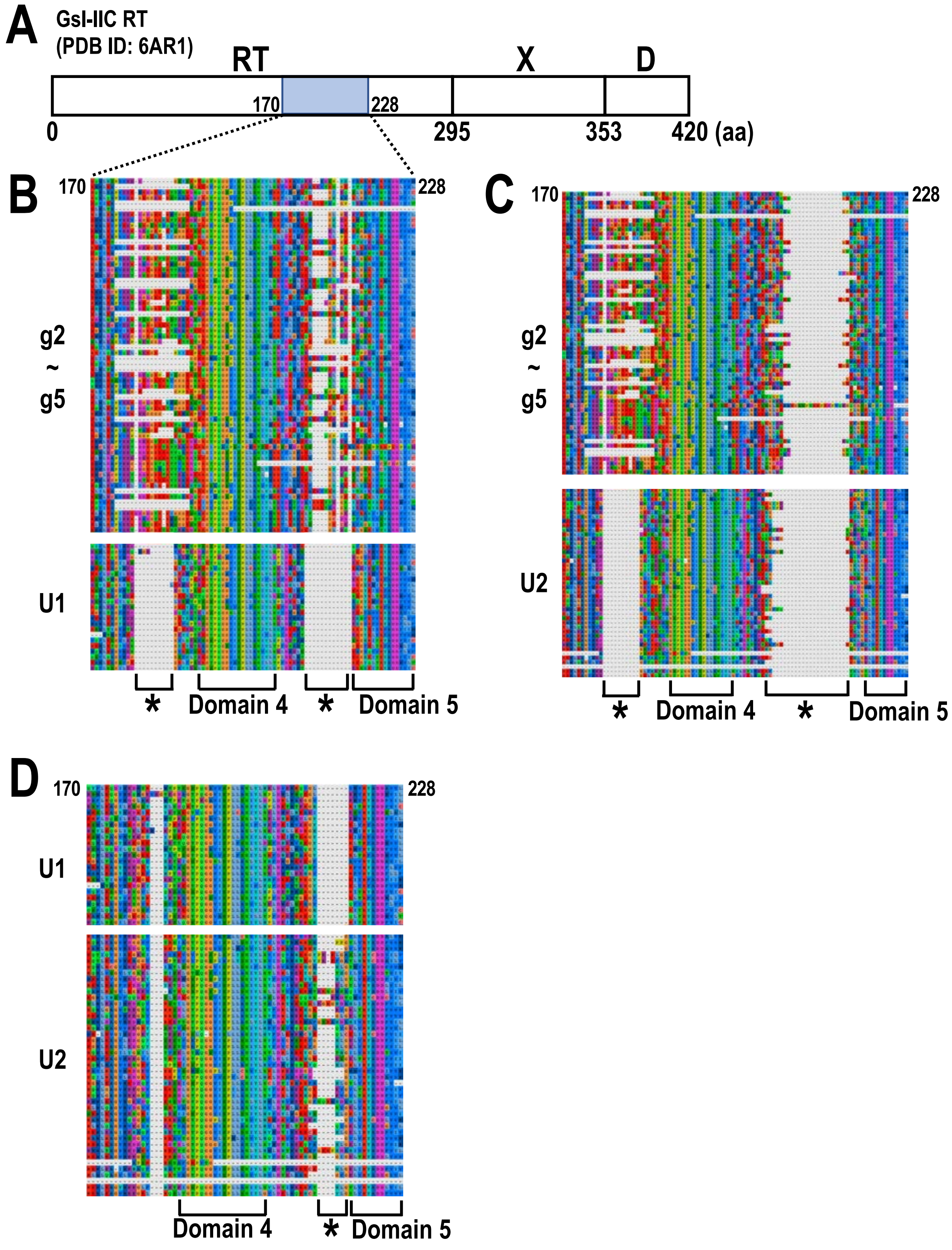


**Supplementary Fig. S6.** Comparison of different amino acid sequence regions among bacterial-g2–g5, U1, and U2 IEPs. (A) Schematic representation of IEPs (domains and amino acid positions are referred to as GsI-IIC RT [Protein Data Bank ID: 6AR1]). RT, reverse transcriptase domain; X, thumb domain; D, DNA-binding domain. (B) Multiple alignment of an amino acid sequence region that differs between g2–g5 and U1 IEPs. (C) Multiple alignment of an amino acid sequence region that differs between g2–g5 and U2 IEPs. (D) Multiple alignment of an amino acid sequence region that differs between U1 and U2 IEPs. Domains 4 and 5 are highly conserved regions among the IEPs (Wang et al. 2011). The asterisks indicate regions that differ significantly between the two IEP types. See also Figure 2 for an overview of the IEPs.

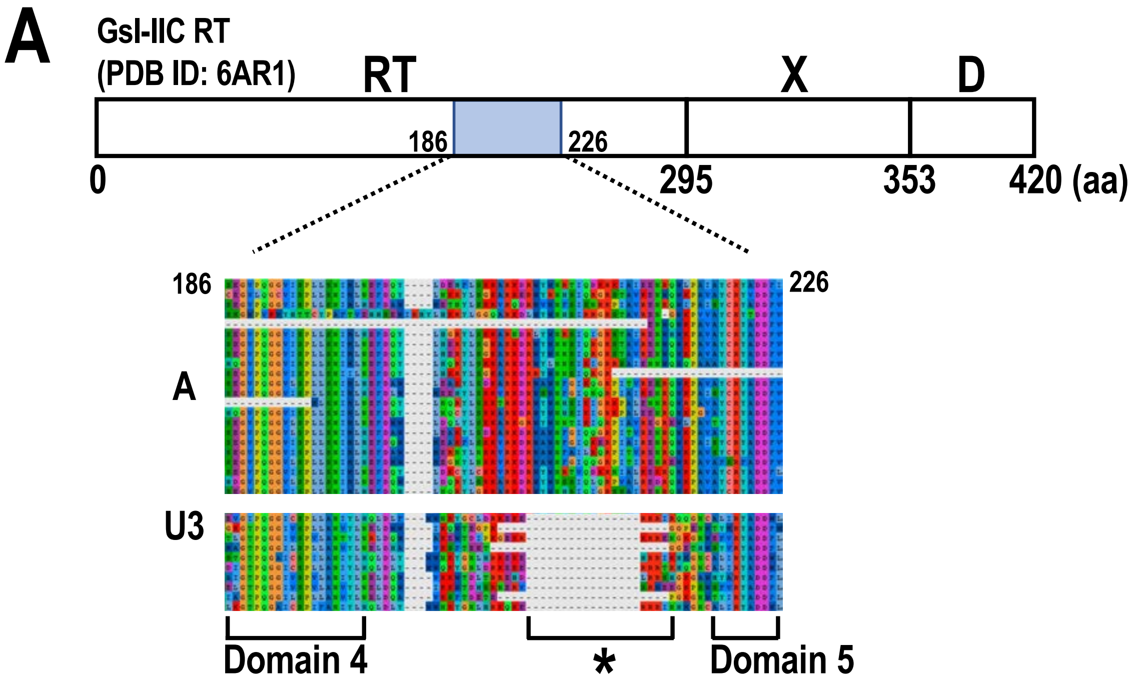


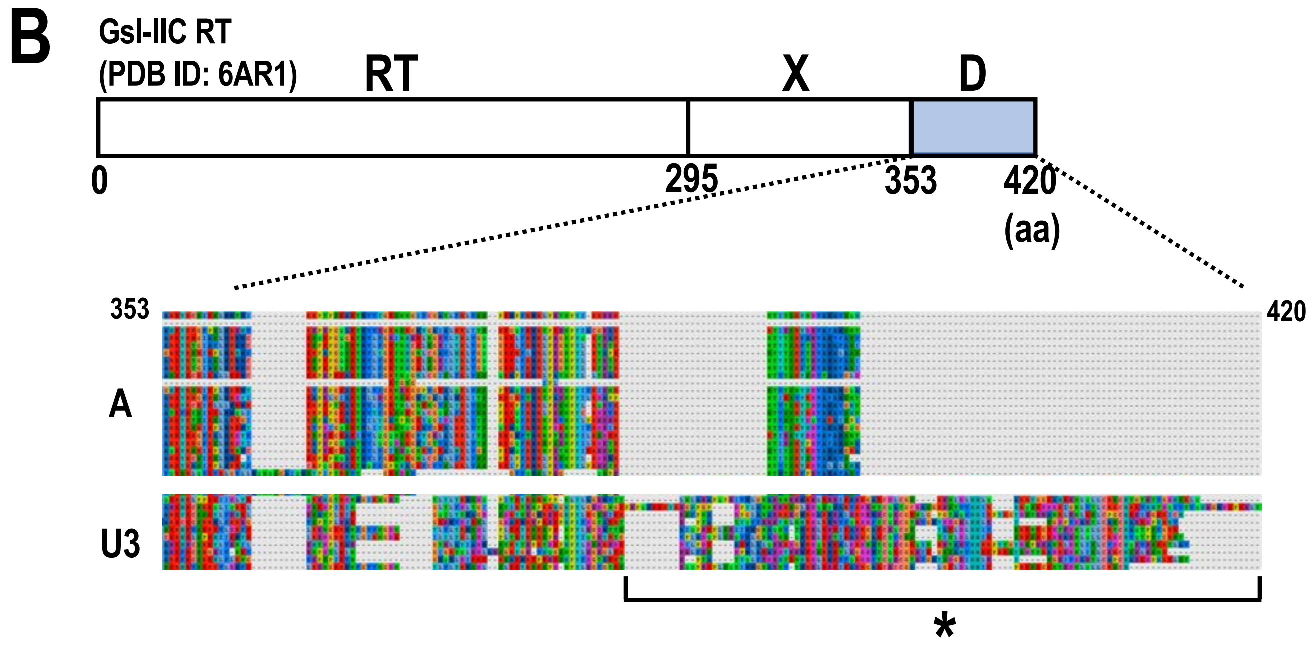


**Supplementary Fig. S7.** Comparison of two different amino acid sequence regions between bacterial-A and U3 IEPs. (A, B) Schematic representation of IEP and multiple alignment of each amino acid sequence region that differs between bacterial-A and U3 IEPs. See the legend to Supplementary Figure S6 for details.

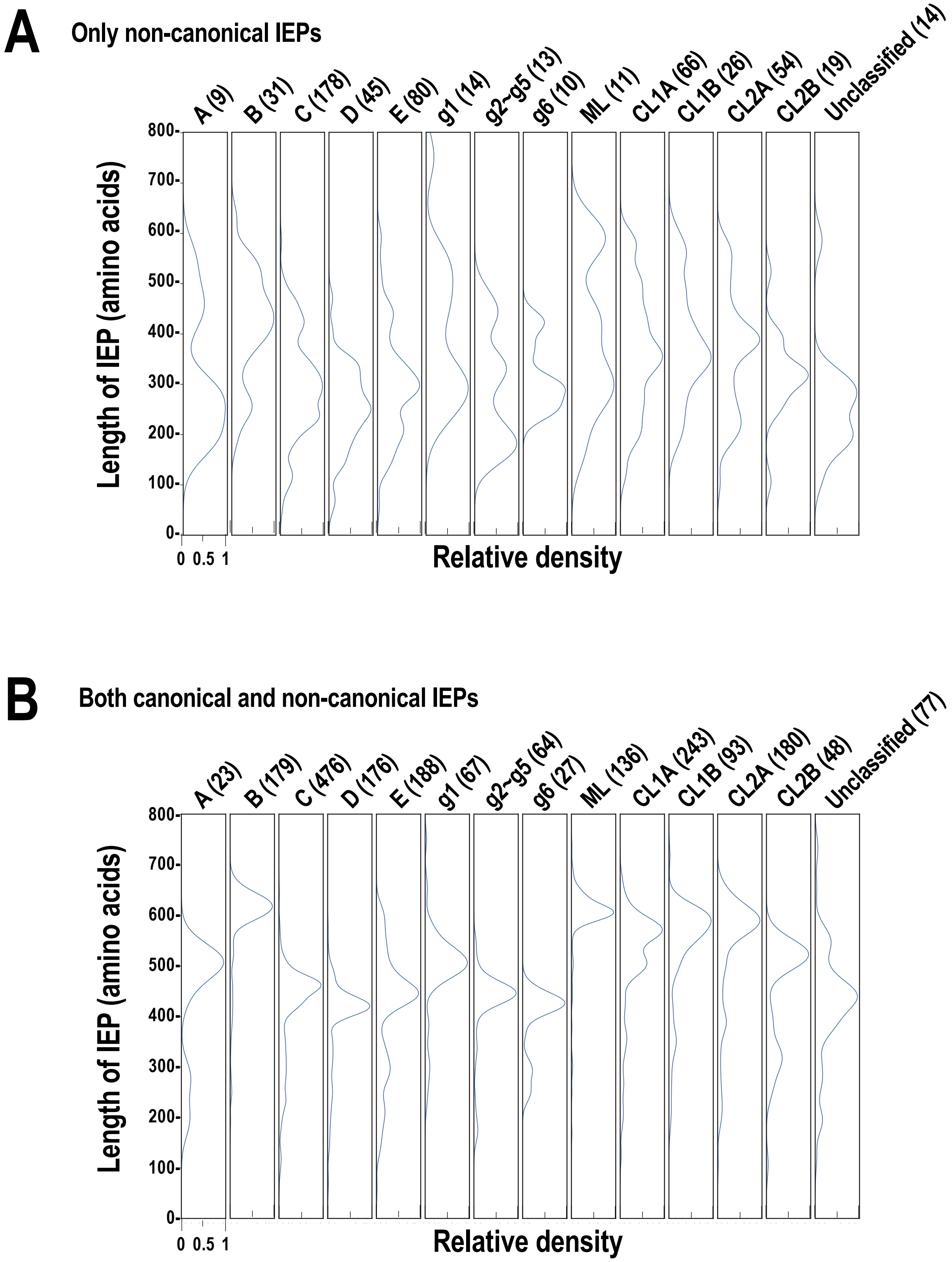


**Supplementary Fig. S8.** Distribution of the amino acid lengths for each type of IEP, including noncanonical types. Distribution of the amino acid length for each IEP type is shown. (A) Distributions of the lengths of ORFs for noncanonical IEPs. (B) Distributions of the lengths of ORFs for both canonical and noncanonical IEPs. The peak relative density was set to 1.0 in each case. See the legend to Figure 2B for details.
