## Supplementary material for "Distinct Expansion of Group II Introns Depends on the Type of Intron-encoded Protein and Genomic Signatures in Prokaryotes": Suppl_Info_II

### Supplementary Information II

**Supplementary Fig. S9 - Supplementary Fig. S14**

#### Distinct Expansion of Group II Introns Depends on the Type of Intron-encoded Protein and Genomic Signatures in Prokaryotes


Masahiro C. Miura,[1,2] Shohei Nagata,[1] Satoshi Tamaki,[1]
Masaru Tomita,[1,2,3] and Akio Kanai[1,2,3*]

[1]Institute for Advanced Biosciences, Keio University, Tsuruoka, Japan
[2]Systems Biology Program, Graduate School of Media and Governance, Keio University, Fujisawa, Japan
[3]Faculty of Environment and Information Studies, Keio University, Fujisawa, Japan

*Corresponding Author

Akio Kanai, PhD
Institute for Advanced Biosciences, Keio University
Tsuruoka, Yamagata, Japan


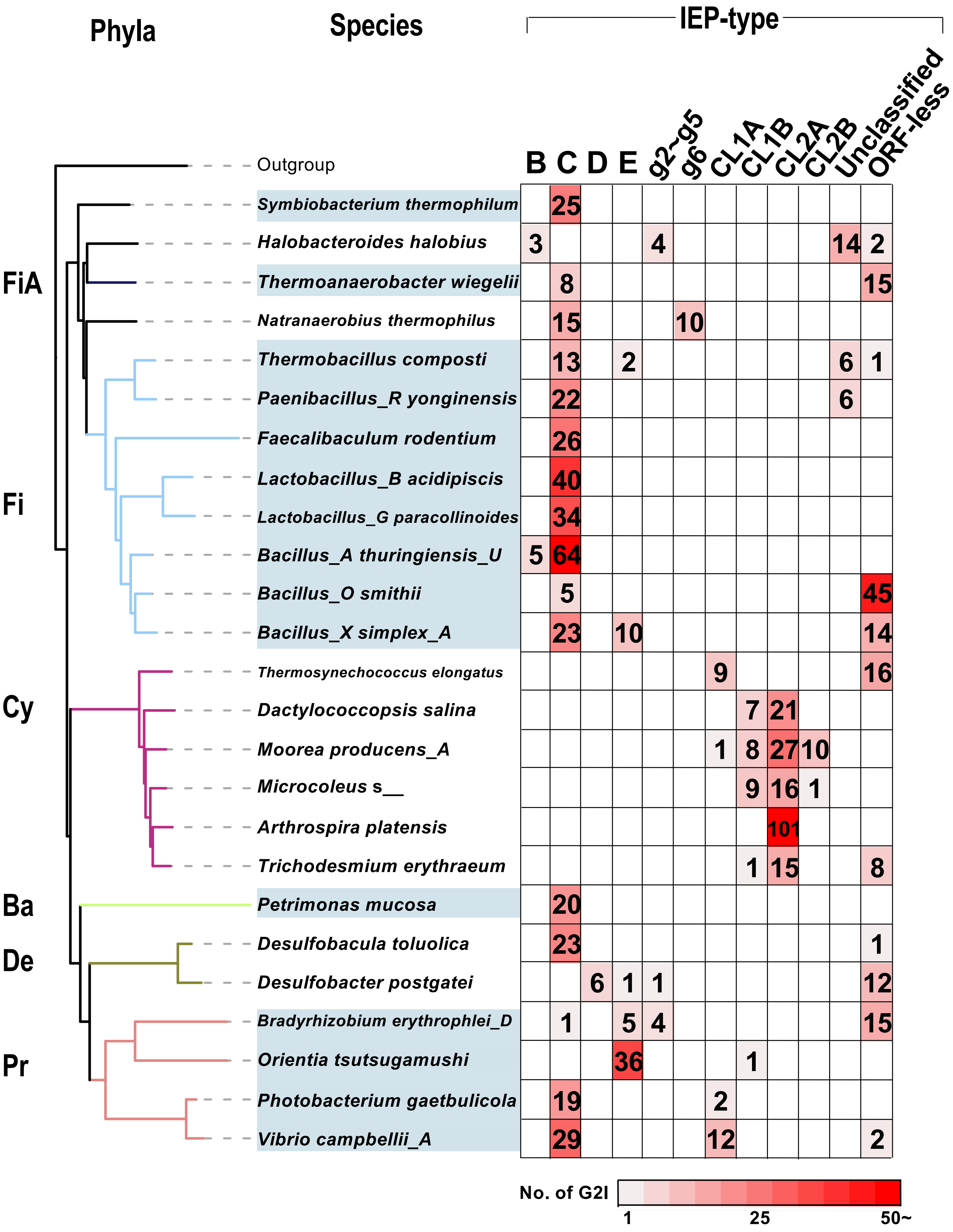


**Supplementary Fig. S9.** Phylogeny of bacteria containing more than 20 G2Is and their types of IEPs. Phylogenetic tree of 25 representative bacteria containing more than 20 G2Is is shown with their types of IEPs. *Candidatus Saccharibacteria* oral taxon TM7x (RefSeq assembly accession: GCF_000803625.1) was used as the outgroup. Heat map indicates the number of G2I(s) per type of IEP. Abbreviations of phyla: FiA, Firmicutes_A; Fi, Firmicutes; Cy, Cyanobacteriota; Ba, Bacteroidota; Pr, Proteobacteria.

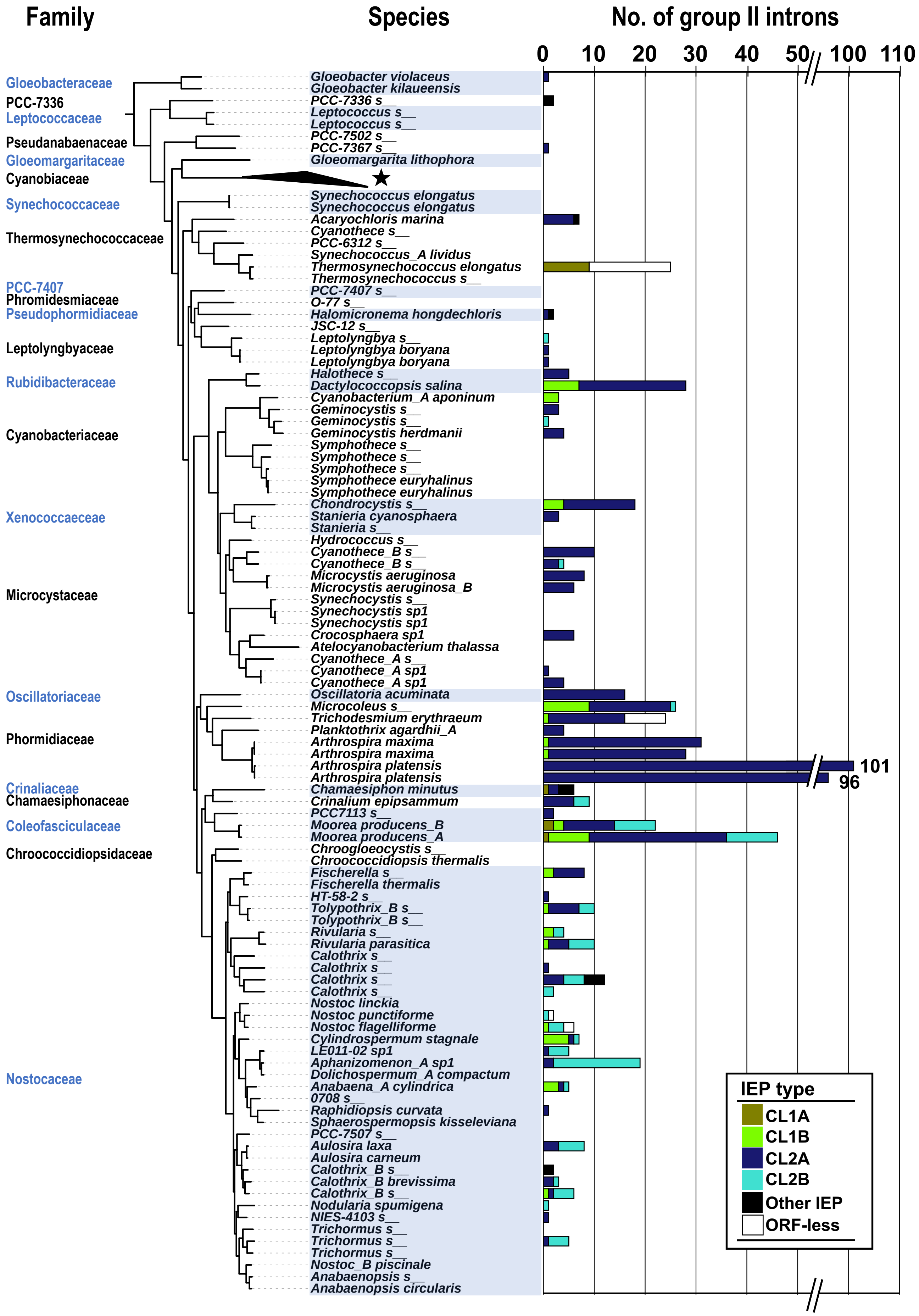


**Supplementary Fig. S10.** Numbers of G2Is and types of IEPs in Cyanobacteriota. The Cyanobacterial phylogenetic tree was constructed from 130 species (see Materials and Methods), shown with their species names. Every other phylum name is written in blue, and the corresponding species are shaded in blue. The numbers of G2Is and the types of IEPs are shown as a colored stacked bar chart in the right box. Because no G2Is were found in the 29 species belonging to the family Cyanobiaceae, the corresponding clade is shown as a triangle and indicated by an asterisk.

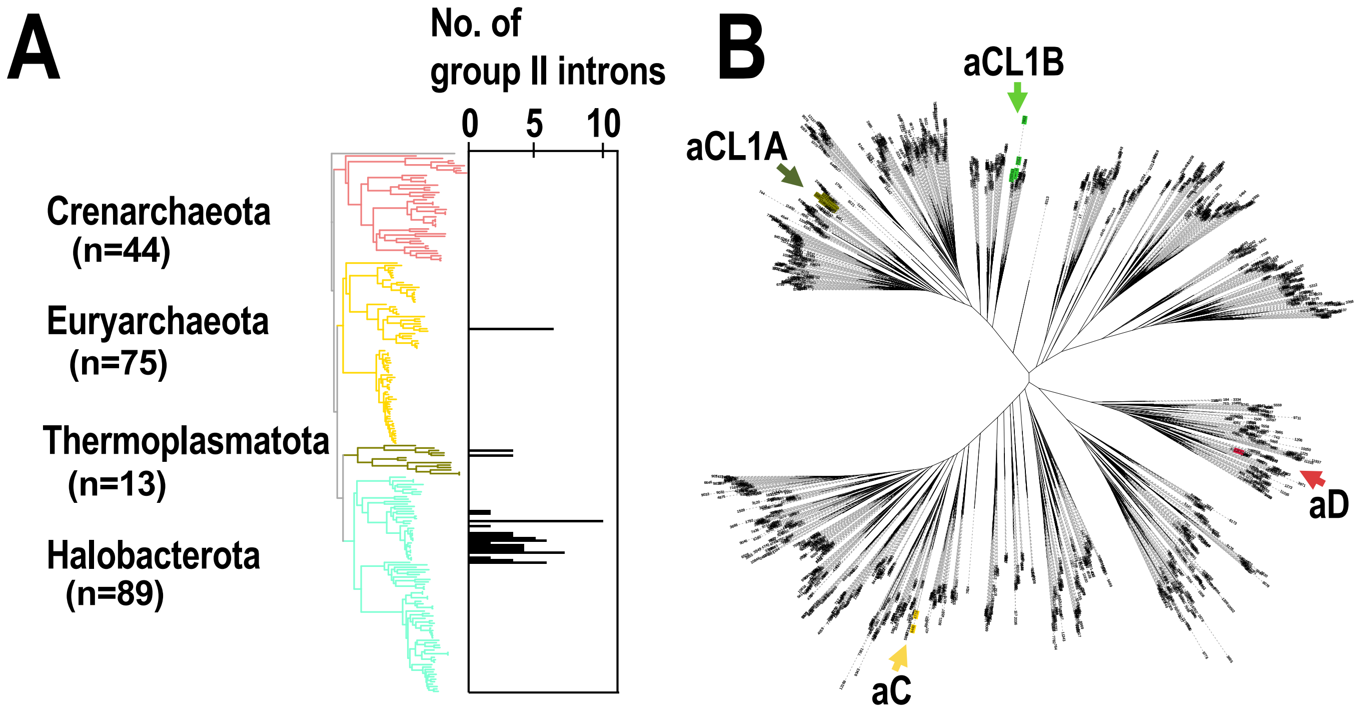


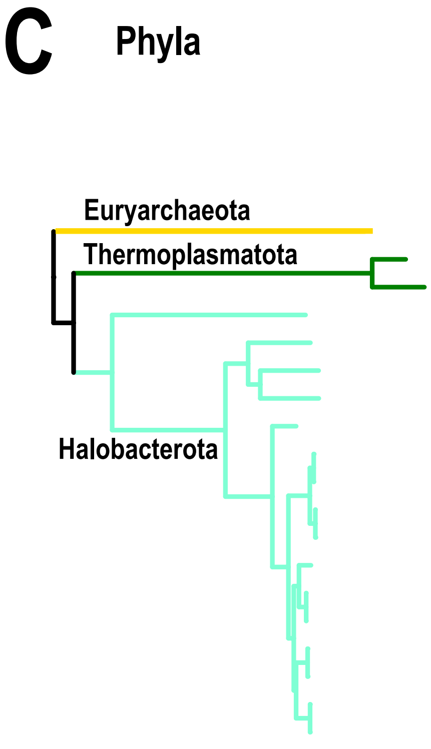


| Species | A | B | C | D | E | g1 | g2~g5 | g6 | ML | CL1A | CL1B | CL2A | CL2B | Unclassified | ORF-less |
|---|---|---|---|---|---|---|---|---|---|---|---|---|---|---|---|
| *Methanobrevibacter_A millerae* | | | | | | | | | | | 7 | | | | |
| *Methanomethylophilus alvus* | | | 1 | | | | | | | | | | | | 1 |
| *Methanomethylophilus* ISO4-H5 | | | 1 | | | | | | | | | | | | 1 |
| *Methanospirillum hungatei* | | | | | | | | | | 6 | | | | | |
| *Methanococcoides burtonii* | | | | | | | | | | 4 | | | | | a |
| *Methanomethylovorans hollandica* | | | | | | | | | | 1 | | | | | |
| *Methanolobus_A psychrophilus* | | | | | | | | | | 1 | | | | | |
| *Methanosarcina sp.* MTP4 | | | | | | | | | | 1 | | | | | |
| *Methanosarcina vacuolata* Z-761 | | | | | | | | | | 2 | | | | | |
| *Methanosarcina vacuolata* Kolksee | | | | | | | | | | 2 | | | | | |
| *Methanosarcina barkeri* MS | | | | | | | | | | 2 | | | | | 1 |
| *Methanosarcina barkeri* CM1 | | | | | | | | | | 3 | | | | | 1 |
| *Methanosarcina lacustris* | | | | 1 | | | | | | | | | | | |
| *Methanosarcina sp2* WWM596 | | | | | | | | | | | | | | | 3 |
| *Methanosarcina sp2* WH1 | | | | | | | | | | | | | | | 3 |
| *Methanosarcina siciliae* C2J | | | | | | | | | | 3 | | | | | |
| *Methanosarcina siciliae* T4/M | | | | | | | | | | 10 | | | | | |
| *Methanosarcina mazei* SarPi | | | | | | | | | | 1 | | | | | |
| *Methanosarcina mazei* S-6 | | | | | | | | | | 2 | | | b | | |

**Supplementary Fig. S11.** G2Is are increased in certain archaeal species. (A) Increases in numbers of G2Is in specific archaeal species. The numbers of G2Is in complete archaeal genomes (222 species) are shown. Archaeal phyla are shown on the left and each corresponding branch on the archaeal phylogenic tree is colored. The numbers in brackets represent the number of species in each phylum. (B) Positions of archaeal IEPs on an unrooted phylogenetic tree of the representative IEP sets (see Figure 2A in details). aCL1A (archaeal CL1A), aCL1B (archaeal CL1B), aC (archaeal-C), and aD (archaeal-D). (C) Distribution of types of IEPs in the archaeal phylogeny. The distribution of the types of IEPs in 19 archaeal species whose genomes contain G2I(s) is shown. Numbers of G2I(s) per IEP type are also shown in each box. a: Analysis of the *Methanococcoides burtonii* genome with our pipeline incorrectly detected four ORF-less G2Is. These were parts of CL1A-type G2Is. b: Analysis of the *Methanosarcina mazei* S-6 genome with our pipeline incorrectly detected one G2I classified as the CL2B type, because the G2I was divided by a transposase. A detailed analysis revealed that it was a CL1A-type G2I.

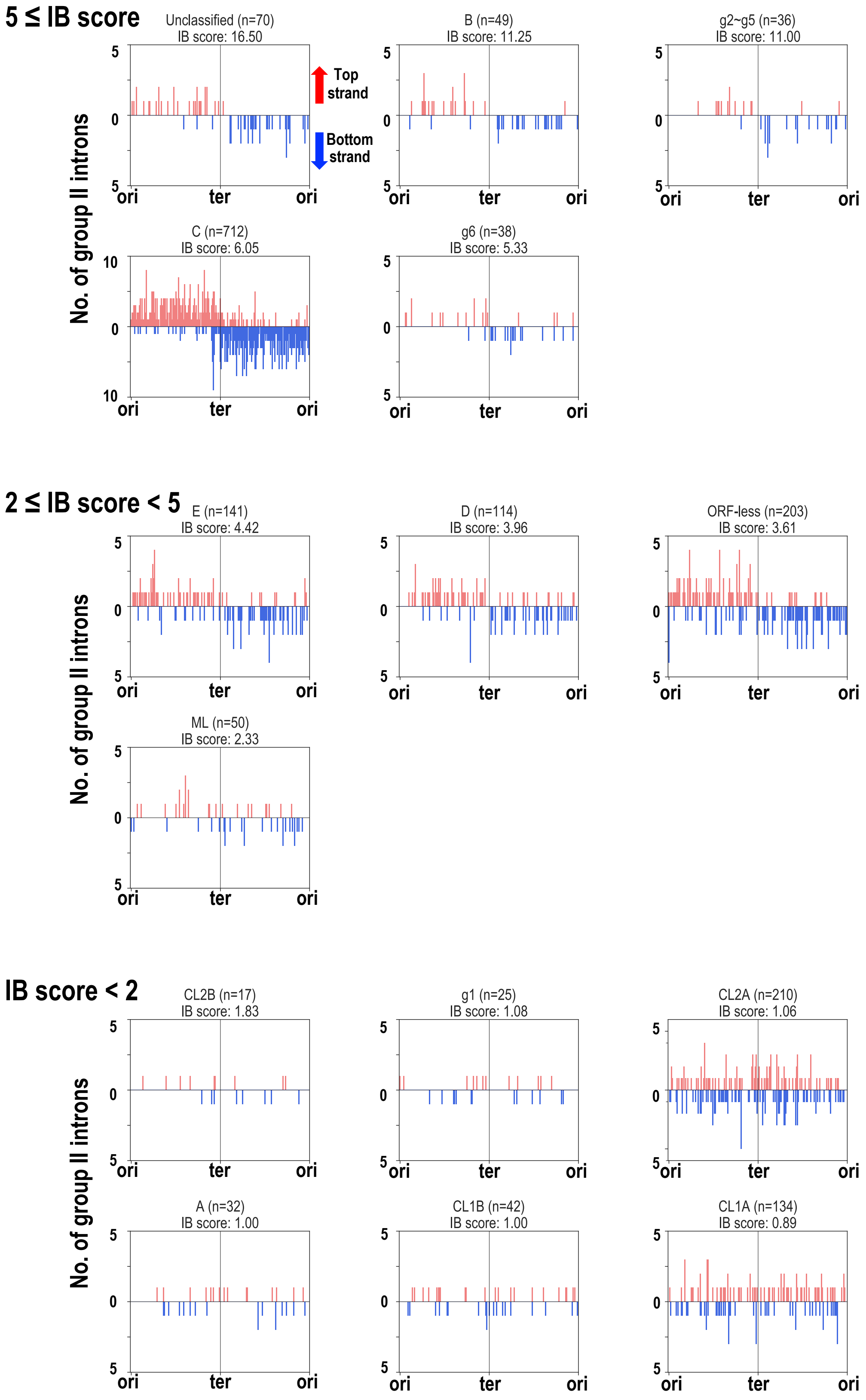


**Supplementary Fig. S12.** Insertion positions of G2Is for each IEP type in the bacterial genomes. Among the representative bacterial species containing G2Is, the insertion positions of the introns in the chromosomes of 349 species for which the position of *oriC* could be obtained from the DoriC database are shown for each IEP type. Positions separated from *ori* by half the chromosome length were defined as *ter*. The right and left arms extending from *ori* to *ter* were each divided into 200 intervals, and the relative insertion positions of the G2Is were determined. In each box, the vertical axis represents the integrated value of the number of G2Is, and the horizontal axis represents the relative position on the chromosome. The numbers of G2Is were plotted in the upper half of each box if they were inserted into the top strand, and in the lower half of the box if they were inserted into the bottom strand. The IEP types and number of G2Is (in brackets) are shown above each box. The Insertion bias (IB) scores were calculated as the ratio of the number of G2Is on the leading strand to the those on the lagging strand and are shown at the top of each box.

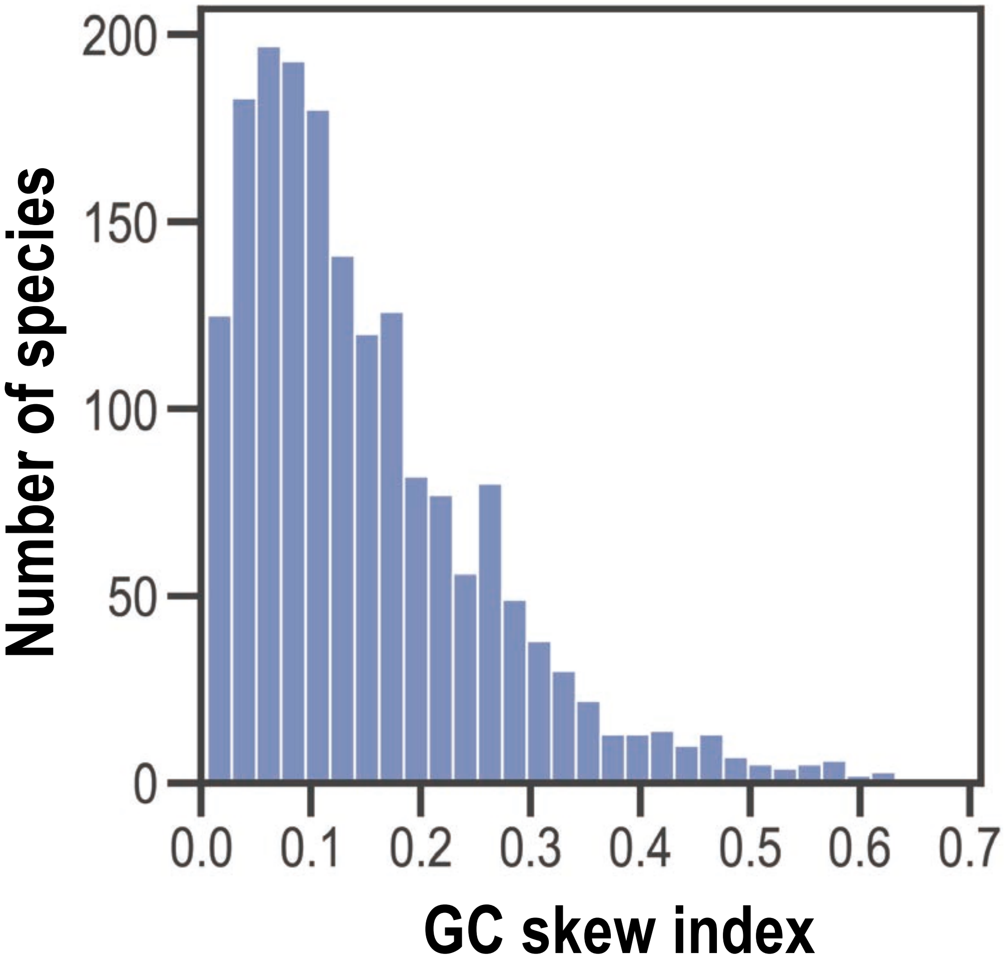


**Supplementary Fig. S13.** Distribution of GC skew indices for bacterial chromosomes (genomes). Distribution of the GC skew indices for the chromosomes of 1,795 representative bacterial species is shown. In this analysis, chromosomes of ≥ 1,000,000 bp were used.

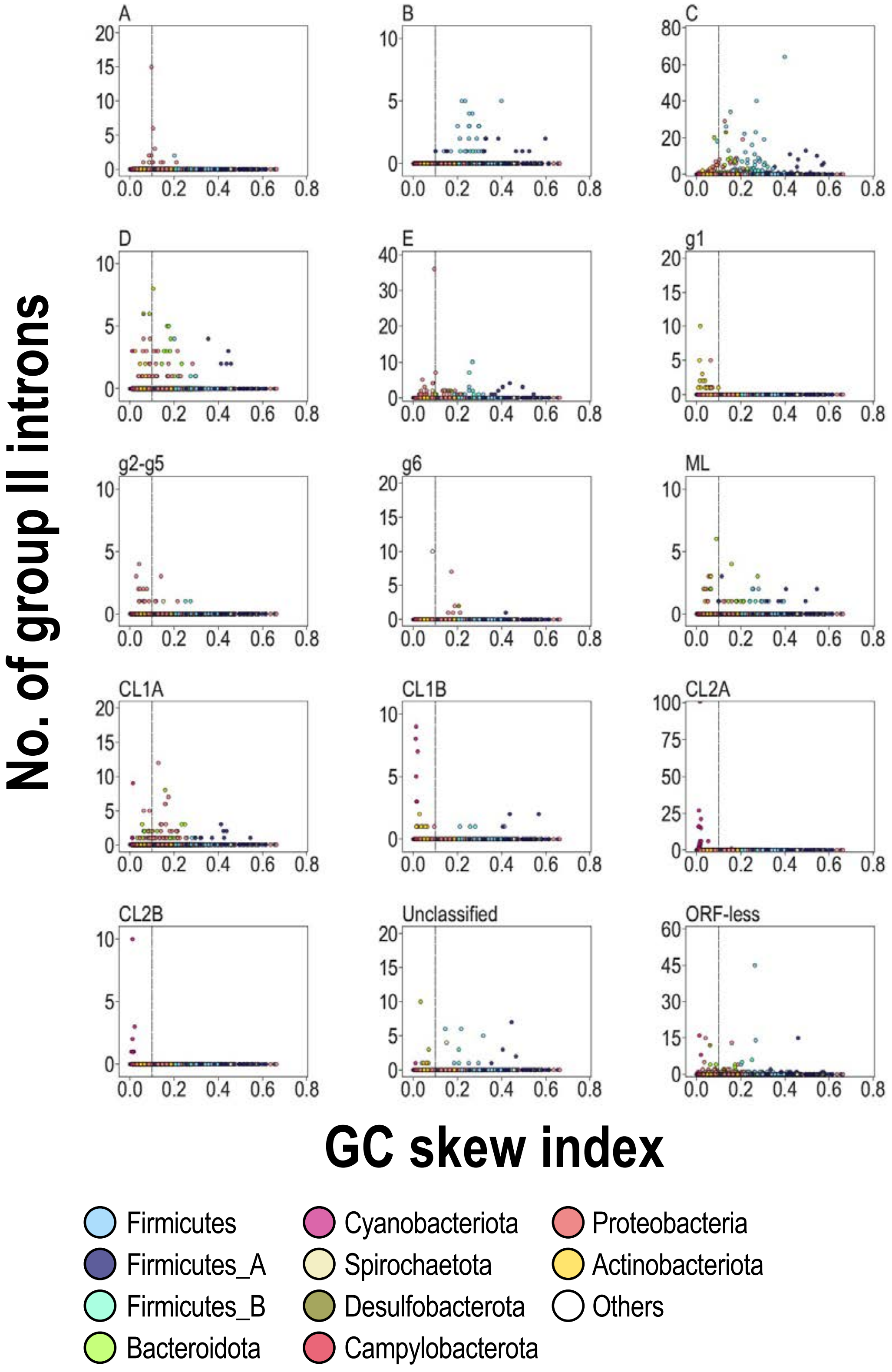


**Supplementary Fig. S14.** Relationship between number of G2Is and GC skew index in bacteria. Relationship between the number of G2Is in the 1,790 representative bacterial species and the GC skew index is shown in a scatterplot for each IEP type. Each dot indicates an individual species, and the color of each dot represents the phylum, as shown at the bottom of the figure. Horizontal axis shows the GC skew index and the vertical axis shows the number of G2Is. The longest chromosomal value for each species was used to calculate the GC skew index.
